# Orthogonal phenotyping from multiplex imaging across scales

**DOI:** 10.64898/2026.11.26.767862

**Authors:** Ali Oğuz Can, Gizem Mansour, Alessandro Palma, Malte Kuehl, Andres Gonzalez Suarez, Kenji Schorpp, Ina Rothenaigner, Kamyar Hadian, Victor G. Puelles, Fabian Theis, Nico Battich

## Abstract

Multiplexed imaging resolves phenotypes across biological scales, from subcellular organelles to multicellular communities. Most representation-learning methods model single scales or combine fine-scale embeddings, conflating signals across levels. An optimal multiscale method would remove inter-scale confounders yielding complementary embeddings — a property we term orthogonal phenotyping, which can be achieved through disentanglement of signatures implicitly arising at each scale. Here we designed scaleVAE specifically for orthogonal phenotyping via a subtractive strategy, a hierarchical variational autoencoder that learns scale-specific representations by conditioning each scale on the preceding one. Through orthogonal phenotyping, scaleVAE resolves multicellular, cellular and subcellular signatures in multiplexed imaging of nuclear membraneless organelles (MLOs) in differentiating human iPSCs, outperforming vision-transformer baselines. With minimal prior knowledge, it separates scale-specific, minimally overlapping phenotypes, revealing how MLO states relate to population structure, differentiation and cell-cycle stage, and showing nonlinear co-variation of polycomb bodies, nuclear speckles and paraspeckles markers across differentiation. In a PathoPlex dataset of mouse crescentic glomerulonephritis, scaleVAE recovered kidney anatomical domains without supervision and resolved pathological subcellular signatures that domain-scale alone did not. This demonstrates that orthogonal phenotyping unlocks complementary biological information that would otherwise remain hidden, establishing it as a powerful framework for automated discovery and a promising foundation for next-generation diagnostic applications.

---

Biological phenotypes are the results of genetic, epigenetic and environmental information flowing between the different biological organizational scales. For example, proliferation and maintenance of stem cells in tissues depend on a highly specialized niche [^1–3^], and differentiated cells adjust their transcript levels by cell intrinsic features, such size, cell cycle stage and the local population context [^4–10^]. In recent years several platforms have emerged for multiplexed imaging of molecular markers allowing spatially resolved omics [^11^], which include iterative indirect immunofluorescence imaging (4i) [^12–14^], PathoPlex [^15^], cell painting [^16–20^], spatial transcriptomics [^21–26^], and spatial proteomics [^27,28^], among others [^29–36^]. These technologies can capture complementary phenotypic aspects at different spatial scales from the multicellular and tissue architecture scales to the cellular and subcellular scales [^37,38^]. However, most artificial intelligence (AI)-based approaches for analysis have focused on learning phenotypic representations at individual scales; either the subcellular scale through the local pixel neighborhood [^12,15,39,40^] or the analysis of full images [^23,41,42^]. While other methods include tailor-made image analysis pipelines, involving object segmentation, and feature engineering for phenotypic quantification [^43,44^]. More recent efforts go beyond the single scale [^45,46^]. However, they consider scales independently [^47–49^] or are bottom-up models that learn embeddings for the finest scale and then combine them to represent higher scales in an additive fashion, thereby lacking mechanisms to ensure extraction of scale specific biological information [^45,50,51^]. Ideally, a multiscale analysis strategy for spatially resolved data should be able to isolate phenotypic aspects operating at each specific scale, while minimizing cross scale interdependency, a property we term *orthogonal phenotyping*.

Here we develop scaleVAE, an approach to achieve orthogonal phenotyping from multiplexed imaging. scaleVAE relies on a multiscale variational autoencoder (VAE) within an unsupervised hierarchical learning architecture to model phenotypic variation from the multicellular towards subcellular scale. Importantly, it allows information flow in the form of a hierarchical conditioning mechanism to prevent the arising of inter-scale confounding factors in a subtractive fashion. This encourages the learning of complementary, non-overlapping representations thereby ensuring orthogonal phenotyping. We validated scaleVAE in two key biological contexts, namely generating a phenotypic atlas of nuclear membraneless organelles (MLOs) across spatial scales during induced pluripotent stem cells (iPSCs) differentiation, and tissue-level extraction of anatomical and signaling features in a kidney injury model, respectively.

Nuclear MLOs are known to regulate transcription[^52–56^], splicing [^57–62^], chromatin states[^63–66^], and participate in the biogenesis ribosomes[^67–70^] and small nuclear ribonucleoproteins [^71–75^]. Using 4i we created a comprehensive dataset to analyze changes in the abundance and spatial distribution of MLOs^76^ in the nucleus, while human iPSCs differentiated into the three germ lineages. scaleVAE was better at disentangling phenotypes across scales than other state-of-the-art network architectures, which allowed us to construct a comprehensive scale-resolved atlas of MLO states during exit from pluripotency. We found that the states of MLOs in a cell depend on differentiation and cell cycle stages, whose relative contributions scaleVAE was able to disaggregate. In addition, MLOs showed specific non-linear patterns of co-occurrences in the nucleus that were associated with specific differentiation stages.

Finally, we applied scaleVAE to multiplexed imaging data from mouse kidney modeling crescentic glomerulonephritis (CGN)[^15^]. This analysis showed that scaleVAE can readily detect tissue anatomical domains and cell states, as well as identifying key phenotypic differences between healthy and injury affected tissues. Specifically, at the subcellular scale, scaleVAE resolved cell states that domain-scale clustering alone did not distinguish, revealing specific pathological signatures in the glomerulus, for example the CGN associated signature of podocytes included a decrease in the expression of DACH1 while p57 remained high. In addition, we also observed that the glomerular region was associated with high levels of the stress marker pJun accompanied by LMP7 and Vimentin.

Together, our analyses across both a developmental (iPSC differentiation) and a disease (kidney injury) context demonstrate the power of scaleVAE to model diverse biological systems and extract orthogonal phenotypes from multiplexed imaging with minimal *a priori* information, enabling the analysis of biological systems across scales to reveal novel insights and potentially accelerate clinical diagnostics.

## Results

### scaleVAE: a hierarchical conditional variational autoencoder

To learn orthogonal phenotypes from multiplexed images, we designed scaleVAE, a hierarchical conditional variational autoencoder (VAE) framework that learns scale-specific latent representations by propagating the learnt latents between scales in a structured manner minimizing the shared information content (**Fig. 1a**). In this context, a scale refers to sampling an image with patches that differ in field-of-view size centering around a shared pixel in the image (**Fig. 1a**). Then, each VAE learns a continuous latent manifold *z_i_* that captures the distribution of its input, which approximates the posterior *q*(*z_i_*|*x_i_*), where *x* is a patch, and *i* is the scale index. To regress out the variation encoded from the previous scale, these latent variables are propagated as conditioning signals to the VAE operating at the next scale. This results in hierarchical phenotype modeling across the defined scales (e.g. multicellular, cellular, and subcellular scales), where the learnt phenotypes capture primarily what was not modeled at the previous scale. Hence, scaleVAE employs a subtractive mechanism to learn scale specific signatures, which contrasts the additive methodology of recent works [^45,50,77^]. In particular, scaleVAE learns a set of condition covariates (*c_i_*) directly from the data, without relying on prior annotations, object segmentations, or other assumptions. The VAEs are implemented as fully convolutional networks that are augmented with multilayer perceptrons for condition encoding and are trained jointly in an end-to-end manner, where gradients are not propagated between VAEs and their interaction is restricted to conditioning. scaleVAE also allows to accommodate inductive biases to guide the extracted information, when possible and desired. This can be done for example, by 1) choosing *a priori* the channels seen at each scale, and 2) restricting the sampled patches to specific objects or regions of the images for object centric modeling. If no such information is available or convenient, scaleVAE will guide the information extraction through conditioning only. We reasoned that this hierarchical modeling strategy would identify the biological scale at which a phenotype emerges in complex multiplexed images and thus allowing quantification of interactions between phenotypes across scales.

**Fig. 1:**
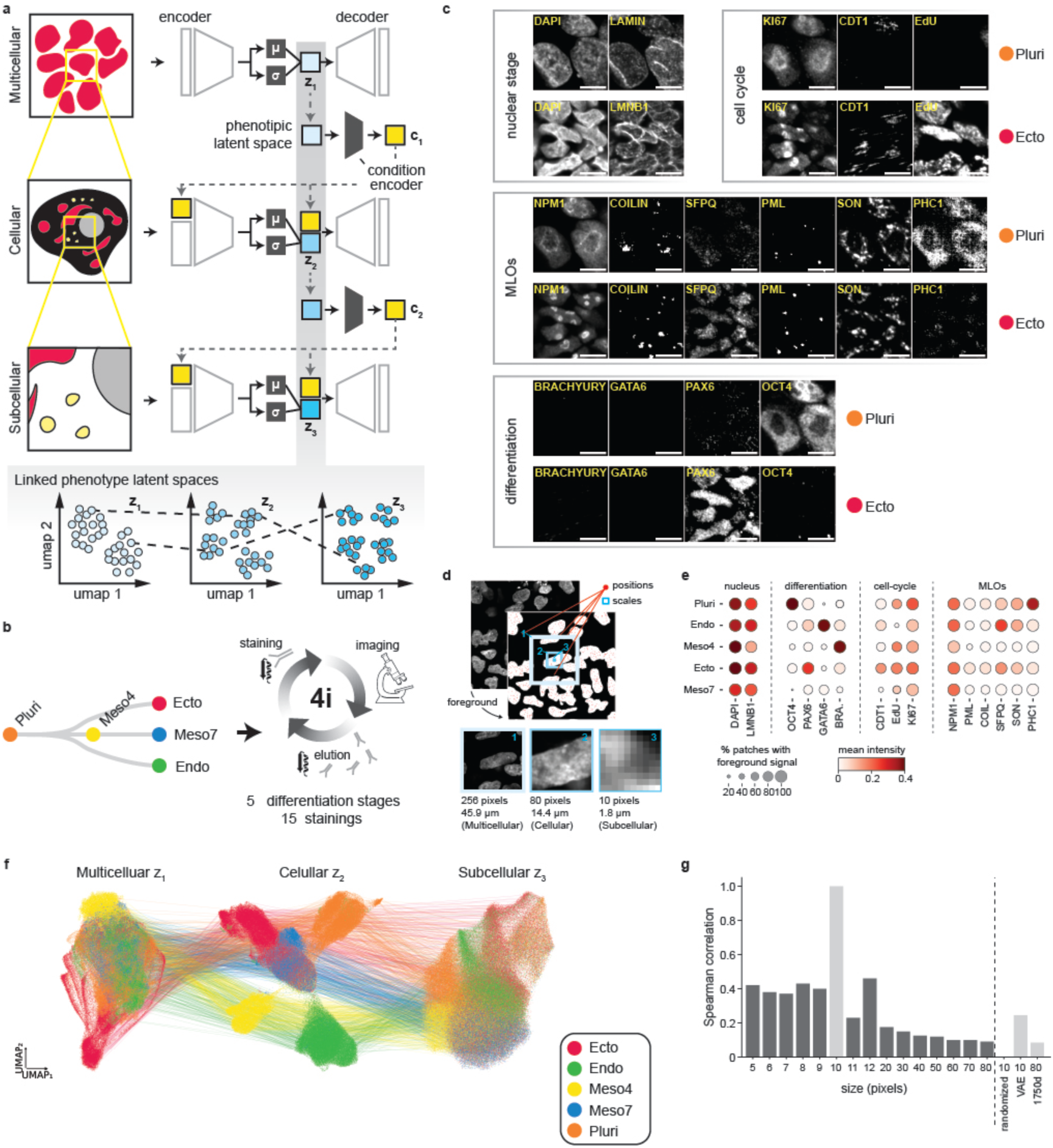
Orthogonal phenotyping from multicellular through subcellular scales on multiplexed images using scaleVAE. **a,** Schematic of scaleVAE architecture for conditional representation learning at each of the three scales. *μ*, mean values; *σ*, standard deviation; *z_i_*, continuous latent manifold comprised of *μ* and *σ*; *c_i_*, condition covariates. **b,** Experiment setup to measure nuclear MLOs during differentiation of human iPSCs (Pluri) into ectoderm (Ecto), mesoderm (Meso) and endoderm (Endo). Cell cultures were stained against 15 markers using 4i after 4 days (Meso4) or 7 days of differentiation (Ecto, Meso7, Endo). As controls, Pluri cells were maintained undifferentiated for 7 days. **c,** Representative images of Pluri and Ecto cultures, stanning for markers of the nucleus, cell cycle stage, differentiation stage, or MLOs are shown. Scale bar, 10 μm. **d,** Schematic of patch sampling for analysis using scaleVAE. **e,** Quantification of mean fluorescence intensity of immunostaining from cell scale patches, dot size reflects the fraction of patches with intensities above background for each staining. The number of patches is 238,091. **f,** UMAP plot of the orthogonal landscapes of phenotypic features corresponding to the latent spaces *z*_1_, *z*_2_ and *z*_3_, lines connect patches of different scales with the same center pixel. **g,** Robustness of scale selection for the subcellular scale, the plot shows the spearman correlation of different patch sizes to a patch of size = 10 pixels. As controls we show a randomized latent space, a variational autoencoder without conditioning (VAE), and a scaleVAE of 80×80 in patch size with a conditional dimensionality of 1,750.

### scaleVAE learns orthogonal phenotypic representations across biological scales in differentiating iPSCs

To study the change in the organization of nuclear MLOs through exit from pluripotency of human iPSCs and test scaleVAE, we created a dataset using 4i [^13,14^] (**Fig. 1b-e**, and **Supplementary Fig.**, **1-3**). Briefly, parallel cultures of human iPSCs were induced to differentiate into ectoderm, mesoderm, and endoderm for one week using standard protocols (**Fig. 1b** and **Supplementary Fig. 3a**, see Methods), while undifferentiated pluripotent cells were also cultured as controls. At indicated time points, 4 days or 7 days after media exchange (**Fig. 1b**), cultures were stained against 15 markers covering nuclear MLOs (SON, PHC1, SFPQ, NPM, COIL and PML), nuclear stage (LMNB1, and DAPI), differentiation stage (OCT4, PAX6, GATA6 and BRACHYURY), and cell cycle stage (KI67, EdU and CDT1) (**Fig. 1c**, and **Supplementary Fig. 3a**). The multiplex images that we obtained with 4i were qualitatively similar to those obtained after standard immunofluorescence (**Supplementary Fig.**, **1** and **2**). Upon visual inspection of the images and direct intensity quantification, we noticed that the abundance of all MLOs change during differentiation hinting at large rewiring of the MLOs states. For example, PML-NBs were more abundant in pluripotent and ectoderm cells than in other differentiation stages, and polycomb bodies and paraspeckles were more abundant in pluripotent and endoderm cells, respectively (**Fig. 1c-e**, and **Supplementary Fig. 3b-c**). To analyze our data using scaleVAE and gain a quantitative understanding of these changes at different scales, we defined square “patches” centered around randomly sampled position within the nucleus of cells with high-quality DAPI signal (**Fig. 1d**, **Supplementary Fig. 4**). The patches had a size of 10 pixels (1.8 µm) for the subcellular scale, 80 pixels (14.4 µm) for the cellular scale, and 256 pixels (46.1 µm) for the multicellular scale (**Fig. 1a** and **d**). We then trained scaleVAE using the sampled patches at each scale (**Supplementary Fig. 5** and **Fig. 1f**), and to ensure phenotypic orthogonality between the different scales we: 1) conditioned the latent representations as described above [^78,79^]; while 2) performed a prior channel selection for the first scale, training the multicellular scale using only the DAPI and LMNB1 markers (**Fig. 1a** and **f**). Since the patch size for the subcellular scale is small (10 pixels), we ensured that the learned phenotypes were robust to small differences in pixel numbers by training scaleVAE with sizes ranging from 5 to 80 pixels at *z*_3_ and quantifying the similarity of the learnt representations relative to patches of size 10 pixels using spearman correlation (see methods). This demonstrated that between the 5 to 12 pixels in size, the phenotypes learned at the subcellular scale were robust (**Fig. 1g**).

To confirm directly the ability of scaleVAE to identify and separate signals arising from different biological scales without prior annotation, we compared the latent representations of the cellular and subcellular scales generated with scaleVAE to those from different state-of-the-art deep learning architectures (**Fig. 2a**). These included an unconditioned VAE (VAE) [^80^], conditional VAE (cVAE) [^79^] that had been conditioned by *a priori* defined cell differentiation culture condition and cell cycle stage, a vision transformer (ViT-MAE) [^81^] and a hierarchical vision transformer (SwinViT) [^51^] (**Fig. 2a**). In a naïve implementation one would choose the markers that define biology at each scale as input channels to ensure orthogonality; however, this is challenging in most applications because the same marker may contain patterns that emerge at different scales, or the scale relevant to a marker might be unknown. Since in our dataset we have well-defined markers of subcellular compartments (MLOs), these can be exploited to measure the ability of scaleVAE to approach optimal scale phenotype disentanglement. Therefore, we also compared the subcellular latent space of scaleVAE trained across all channels (scaleVAE), to that of a VAE trained using only MLO markers (MLOs only), as well as scaleVAE trained using only MLO markers in the subcellular scale (MLO only + scaleVAE) (**Fig. 2a**). We reasoned that if scale disentanglement of phenotypes was achieved, markers of MLOs should preferentially drive the formation of the latent space at the subcellular scale (*z*_3_). To test this, we devised two approaches (**Fig. 2b-c**) that aim to quantify the clustering as well as the global manifold structure of the learnt latent representations.

**Fig. 2:**
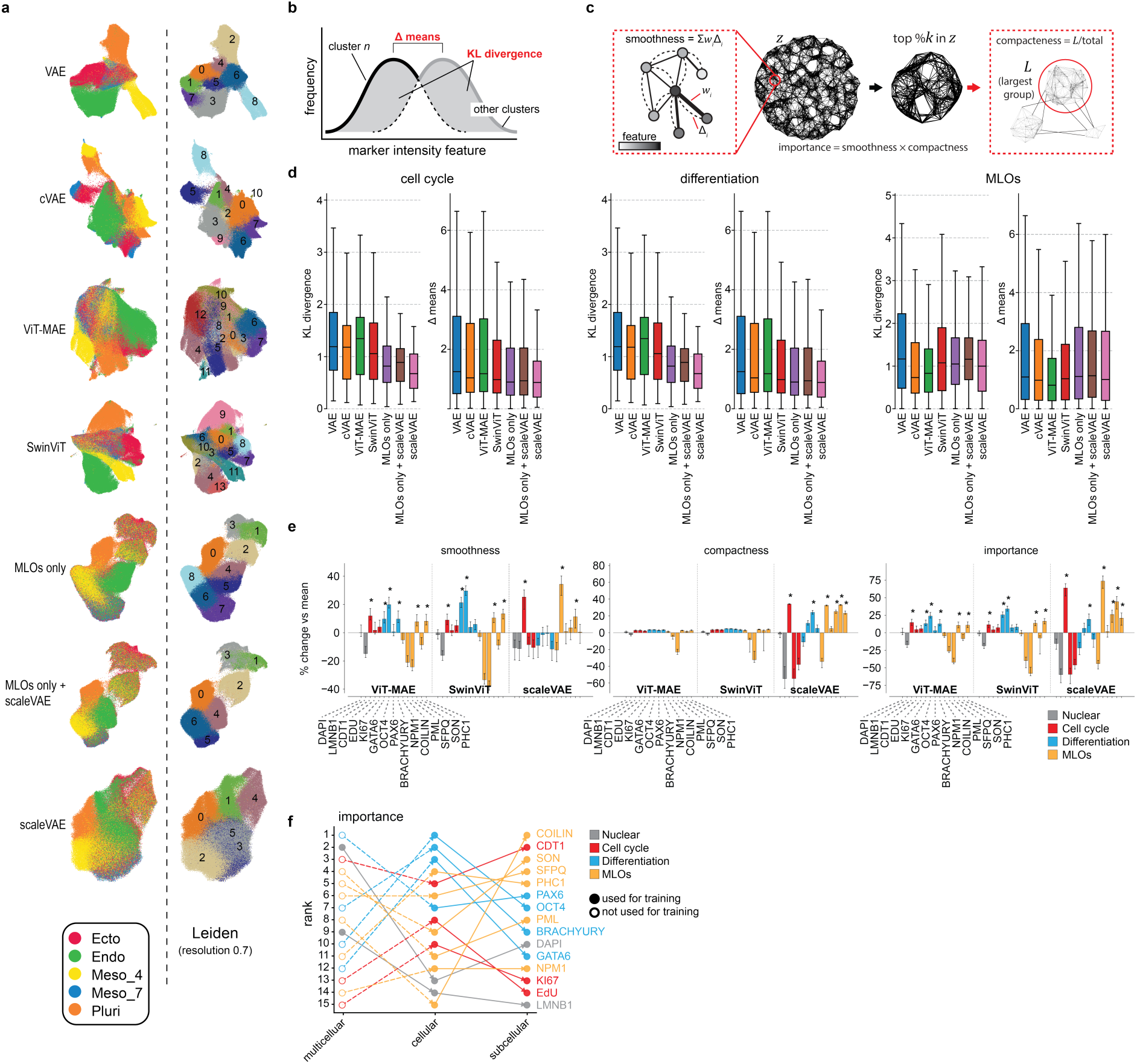
Orthogonal phenotyping across biological scales using scaleVAE. **a,** UMAP projections of phenotypic latent spaces of patches at the subcellular scale (*z*_3_) color-coded by culture condition (left), and Leiden partition (right). Models are; VAE: unconditioned variation autoencoder, cVAE: variational autoencoder conditioned with prior knowledge (cell cycle and differentiation, see Methods), ViT-MAE: a vision transformer masked autoencoder, SwinViT: Swin Transformer, MLOs Only: VAE trained only with MLO markers, MLOs Only + scaleVAE: scaleVAE trained with only MLO markers at subcellular scale, and scaleVAE. **b,** schematics of the differential feature analysis using differences of means and the corresponding Kullback-Leibler (KL) divergence. **c,** Schematics showing the computation of the smoothness, compactness and importance values. **d,** Differences in means and KL divergences (as in **b**) of clusters for cell cycle, differentiation and MLOs features. **e,** Relative smoothness, compactness and importance of the subcellular scale normalized to the mean values across all markers for ViT-MAE, SwinViT and scaleVAE models. *p* values are one-sided Mann-Whitney U tests corrected by Benjamini-Hochberg on 20 bootstraps of 30,000 samplings without replacement, * *p*<0.05 and positive effect size > 5%. **f,** The ranking of markers across scales based on importance is shown for scaleVAE. Rank 1 means the most important stain for defining the latent manifold.

We first performed a one-vs-all differential feature analysis on the Leiden partition of *z*_3_ comparing the means and the corresponding Kullback-Leibler (KL) divergences of features calculated from the intensities of markers grouped into cell cycle, differentiation, and MLO categories (**Fig. 2a** and **b, Supplementary Fig. 6**, see Methods). We expect that markers driving the partition will show relatively high differences in feature magnitudes between the obtained clusters. More specifically for the subcellular scale, if we achieve orthogonal phenotyping, we expect features from cell cycle and differentiation markers to be similar between clusters, while MLO marker features to differ. Note that the extent of achievable disentanglement of the scales is inherently constrained by shared biological information between the scales. We found that scaleVAE outperformed other methods, keeping the difference in features of cell cycle and differentiation low between clusters, while maintaining a high differences between MLO features, without the need for manual specification of conditional covariates, unlike the cVAE, SwinViT and ViT-MAE architectures (**Fig. 2d** and **Supplementary Fig. 6**). Indeed, scaleVAE performed at least as well as the network architectures trained only with MLO markers in the subcellular scale (MLOs Only, and MLOs Only + ScaleVAE) (**Fig. 2d** and **Supplementary Fig. 6**).

Second, we further confirmed the ability of scaleVAE to identify appropriate signatures for a given scale without needing to cluster the latent spaces. To this extent, we introduced *smoothness*, *compactness* and *importance* values using the Rayleigh quotient on the Laplacian graph and the cardinality of the largest connected component of the neighborhood graph (i.e. the top 5% samples with respect to the mean intensities) (**Fig. 2c**, **Supplementary Fig. 7**, see Methods) [^82^]. The smoothness metric measures the gradualism in the changes of a marker over the entire geometry of the latent space, while compactness measures whether the highest intensities are group together (**Supplementary Fig. 7a**). Therefore, the derived importance value can be seen as a proxy for the impact that a marker has in organizing the latent space, i.e. driving a smooth and compact organization (**Supplementary Fig. 7a**). We compared the relative smoothness, compactness and importance values for the mean intensity of all markers at the subcellular scale for ViT-MAE, SwinViT and scaleVAE models. A positive relative value indicates markers with larger than the mean values across all markers for a given model. We found that SFPQ and PHC1 (two out of six MLO markers) were significantly increased over the mean in smoothness and importance for the ViT-MAE and SwinViT models. While for scaleVAE, two (COILIN and SON) out of six were significantly increased in smoothness, and four (COILIN, SFPQ, SON, and PHC1) were significantly increased in compactness and importance (**Fig. 2e**). These findings were robust to changes in the parameters for value computation **(Supplementary Fig. 7b-e**), and demonstrated that scaleVAE outperforms other architectures disentangling scale phenotypes. Furthermore, when looking at the rankings of importance across the different scales, we found that, in scaleVAE, among the top five most important markers organizing the latent space of the subcellular scale (*z_3_*), four were MLO-specific (COILIN, SON, SFPQ and PHC1), while in the cellular scale only PHC1 was found within the top five (**Fig. 2f**). This may be explained by the high expression for the PHC1 in the pluripotent state. In contrast, SwinViT and ViT-MAE had only two (PHC1 and SFPQ) and one (PHC1) MLO markers that ranked in the top five organizers of the subcellular latent space, respectively (**Supplementary Fig. 7e**). Using covariance as a proxy for linear orthogonality, we compared each network’s subcellular and cellular latent spaces. scaleVAE’s subcellular space differed from its cellular space by 92%, compared with 77% for the best non-scaleVAE approach (**Supplementary Fig. 7f**). We next tested the conditional strength required to achieve orthogonal phenotyping, and found that above a condition dimensionality of 70, scaleVAE remained robust (**Supplementary Fig. 7g**). This last result shows the importance of choosing a sufficiently high condition dimensionality.

We next tested the conditional strength required to achieve orthogonal phenotyping, and found that above a condition dimensionality of 70, scaleVAE remained robust (**Supplementary Fig. 7g**). This last result shows the importance of choosing a sufficiently high condition dimensionality. Importantly, scaleVAE automatically assigns distinct biological signatures to different latent spaces, effectively disentangling the scales and achieving orthogonal phenotyping of complex multiplexed imaging datasets.

### At the multicellular scale cell density is coupled to cell-cycle and MLO marker levels

To determine whether population-level context shapes nuclear phenotypes, we interrogated the multicellular scale latent space (z1) by Leiden clustering and computing the mean foreground fraction per cluster, as a proxy for local cell density (see Methods; **Fig. 1a** and **d**). We run scaleVAE independently in two biological replicates (Rep1 and Rep2) and analyzed the resulting clusters together. Both replicates presented distinct population types spanning a wide range of densities made from clusters that had different culture condition compositions (**Extended Data Fig. 1a**). Ranking clusters by descending foreground fraction showed a visible gradient from higher to lower cell-cycle and MLO marker intensities correlating to the cell density (**Extended Data Fig. 1b**). Although most clusters contained a mix of germ layer and pluripotency populations (**Extended Data Fig. 1b**, **Supplementary Data 1**; **Supplementary Fig. 8a**), denser clusters (e.g. clusters 14 Rep2, 9 Rep2, and 18 Rep1) were predominantly pluripotent- or ectoderm-dominant, clusters of intermediate density (e.g. clusters 3 Rep1, 17 Rep1, 13 Rep1) were predominantly endoderm-dominant, and the sparsest clusters (e.g. clusters 12 Rep2, 10 Rep1, 15 Rep2) were mostly mesoderm-dominant (**Extended Data Fig. 1b and c**). The average foreground fraction across clusters correlated with average cell-cycle marker intensity (|ρ| = 0.65, p = 2.6 × 10^−5^) and average MLO marker intensity (|ρ| = 0.69, p = 5.2 × 10^−6^) when pooling clusters from both replicates (**Supplementary Fig. 8b**). All individual MLO and cell cycle marker intensities strongly correlated with the foreground fraction (average |ρ| range = 0.50–0.66), except SFPQ (average |ρ| = 0.25; **Extended Data Fig. 1d**). These results demonstrate that at the multicellular scale, the cell cycle dynamics and the levels of MLOs are tightly coupled to the population structure. This observed density-dependent pattern agrees with previous work showing that cell density modulates proliferation and differentiation [^9,10^].

**Extended Data Fig. 1:**
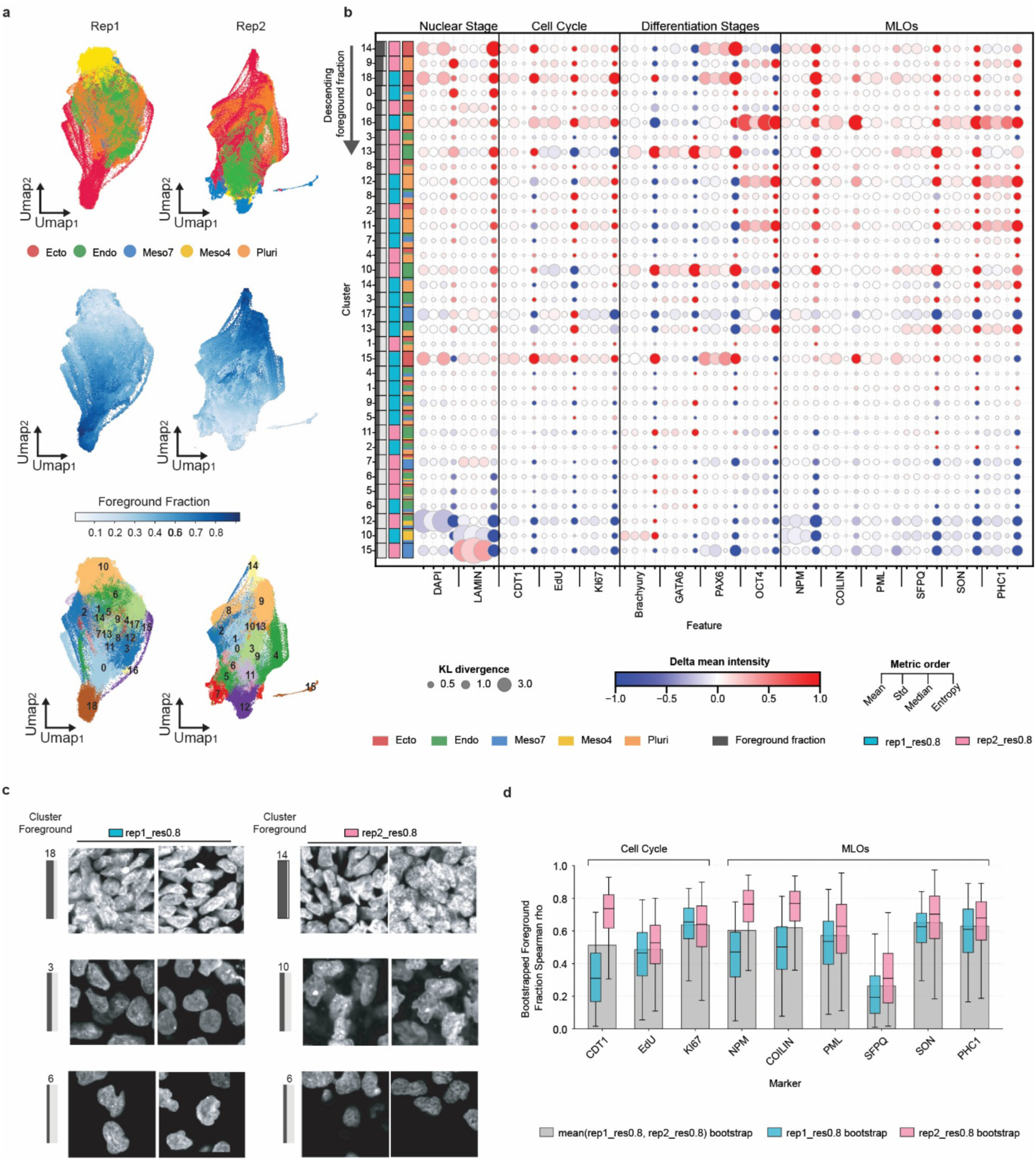
The phenotypic landscape of the multicellular latent space (z_1_). **a,** UMAP projections of the multicellular scale latent space (z_1_) for two independent differentiation experiments (Rep1, Rep2), colored by culture condition (top), foreground fraction (middle), and Leiden cluster assignment (bottom). **b,** Dot plot of differential feature analysis across Leiden clusters of z_1_ (Rep1 and Rep2 combined; 19 and 16 clusters, respectively), ranked by descending mean foreground fraction, first property bar. Replicate identity is indicated by the second property bar. The fraction of patches from each germ layer/pluripotency culture condition contributing to a cluster is indicated by the third property bar. Columns in the dot plot represent mean, std., median and entropy of intensity for nuclear-state, cell-cycle, differentiation and MLO markers. Dot color indicates delta mean intensity; dot size indicates the corresponding KL divergence (see Methods). **c,** Representative images of multicellular patches of three representative Leiden clusters spanning high to low foreground fraction for each replicate; two example patches are shown per cluster. The foreground fraction of each cluster is indicated by the shaded property bar at the left. **d,** Bootstrapped (5,000 iterations, sampling clusters with replacement; see Methods) absolute Spearman correlation between per-cluster foreground fraction and cell-cycle or MLO marker intensity, computed independently for Rep1 and Rep2. Whiskers in box plots, 2.5th–97.5th percentile.

### Cellular scale phenotyping resolves cell-cycle and differentiation effects on the organization of nuclear MLOs

The mixed germ layer composition of clusters in the multicellular scale was well suited to studying the impact of the population density on nuclear organization, yet it did not allow us to resolve the influence of the cell cycle and differentiation stages (**Extended Data Fig. 1a**). We therefore examined the cellular scale latent space (z_2_), whose UMAP revealed a clear structure organized by both differentiation and cell-cycle markers (**Fig. 3a**). We reasoned that this structure in z_2_ should allow us to assess how cell-cycle progression and differentiation stage jointly shape MLO organization. To do this, we first partitioned z_2_ into Leiden clusters, computed features from the intensity of markers in the cell scale patches (mean, standard deviation, median and entropy, see Methods), and then annotated the clusters by cell-cycle (**Supplementary Fig. 9a, b**) and differentiation stages (**Supplementary Fig. 10a**; **Supplementary Data 1**). Clusters with zero or very low pixel intensities in all markers, and clusters corresponding to M-phase, were removed from the analysis (see Methods). To test whether the phenotypic organization of nuclear MLOs was reproducible for the remaining clusters in Rep 1 and Rep 2, we compared them by cosine similarity of the differential feature analysis of the mean intensities (z-score delta mean intensities and its KL divergence; **Fig. 3b**): clusters of the same differentiation stage showed high cross-replicate similarity (**Fig. 3b**; **Supplementary Fig. 10b**). However, this correspondence across all markers could in principle reflect shared differentiation or cell-cycle marker levels (e.g., shared high GATA6 in endoderm clusters) rather than a genuine correspondence in MLO organization. After restricting the similarity comparison to the six MLO markers only, we observed similarly high cross-replicate correspondence (**Supplementary Fig. 10c**), indicating that MLO organization itself is reproducible across replicates.

**Fig. 3:**
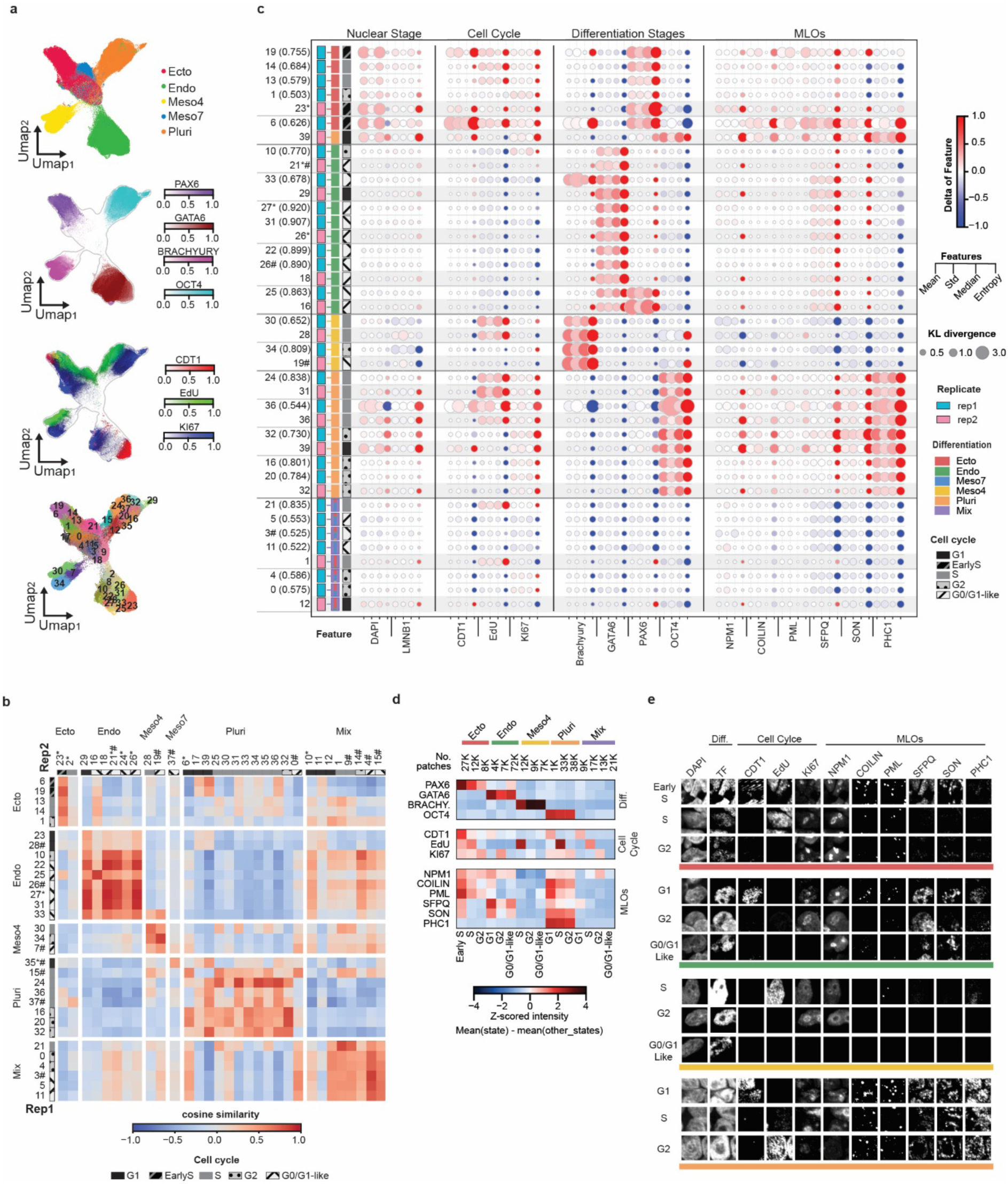
The phenotypic landscape of the cellular latent space (z_2_). **a,** UMAP projections of the cellular scale latent space z_2_ (Rep 1), colored by culture condition, differentiation marker intensity, cell-cycle marker intensity, and Leiden cluster assignment, top to bottom, respectively. **b,** Cosine similarity between cellular scale Leiden clusters of Rep 1 (rows) and Rep 2 (columns; see Methods). Clusters are grouped and ordered by differentiation stage; Mix denotes clusters with mixed culture condition composition; shown as a purple-framed segment whose color proportions reflect that composition. Bars at the left and top indicate the assigned cell-cycle stage. **c,** Dot plot of differential feature analysis for the cellular scale for Rep 1 and matched Rep 2 clusters (see Methods), sorted by differentiation stage and cell-cycle stage. Dot color indicates delta of feature and dot size indicates KL divergence. Side bars indicate group, replicate, differentiation and cell-cycle. # denotes a low-DAPI cluster; * denotes an ambiguous cell-cycle assignment**. d,** Effect size of differentiation, cell-cycle and MLO marker mean intensity across for grouped clusters according to equivalent differentiation stage and cell-cycle stage. The effect size is the patch-weighted, z-scored differential of delta mean intensity of each state (see Methods). Only states represented among the matched clusters in c are shown. The number of contributing patches is indicated above each column. **e,** Representative images of individual cellular scale patches for different cell-cycle stages and differentiation stages. TF indicates the transcription factor marker characteristic of that differentiation stage (PAX6 for Ecto, GATA6 for Endo, Brachyury for Meso4, OCT4 for Pluri).

To construct the phenotypic landscape of the cellular scale and resolve the interaction between individual cell-cycle-by-differentiation stages with specific MLO phenotypes, we paired Rep 1 clusters with their best-matching Rep 2 cluster (cosine similarity ≥ 0.5) and explored the differential feature analysis from the four features mentioned above (**Fig. 2b** and **Fig. 3c**). We sorted the paired clusters by the differentiation stages (middle bars in **Fig. 3c**) along with their quantified cell cycle states (black and white bars in **Fig. 3c** and **Supplementary Fig. 9**) and noticed that the differentiation stage of the clusters correlated to distinct MLO feature profiles (e.g. PHC1 features were high in the pluripotent stage, SFPQ features were high in endoderm, and mesoderm had overall low MLO feature values). In addition, within a given differentiation stage some MLOs showed differential feature values corresponding to specific cell cycle states (e.g., features of SON in ectoderm early S cluster 6 vs S cluster 14), demonstrating a strong statistical interaction between differentiation stage and cell-cycle stage impacting MLO phenotypes. To better quantify this cell-cycle-by-differentiation stage impact on MLOs, we matched clusters with equivalent combined states and run an effect size analysis (see Methods; **Fig. 3d**, **Supplementary Data 2**).

The intensity of differentiation markers closely tracked assigned differentiation stage identity across the matched cluster sets (**Fig. 3c** and **d**). Brachyury was most enriched in early mesoderm (Meso4) clusters (effect size 3.7 to 3.8), GATA6 in endoderm (1.7 to 2.8), PAX6 in ectoderm (1.8 to 3.0), and OCT4 in pluripotent clusters (2.0 to 2.2), consistent with a correct assignment of differentiation stage identity. Likewise, cell-cycle marker intensity followed the pattern expected from the cell-cycle scoring criteria (**Supplementary Fig. 9**). CDT1 was the dominant marker in G1 states (0.19 to 0.82), EdU in S-phase states (1.15 to 2.65), and KI67 in G2 states (0.45 to 0.96 in most groups), while all three markers were low in G0/G1-like states. We found that cell-cycle markers varied within the differentiation stages, for example CDT1 had the heist level in ectoderm and EdU in pluripotent cells (**Fig. 3c** and **d**). This differentiation-stage-specific variation in cell-cycle markers is consistent with the established coupling between cell-cycle state and cell fate in pluripotent stem cells. Human PSCs characteristically exhibit an abbreviated G1 phase and spend a large proportion of the cell cycle in S phase, whereas lineage commitment is accompanied by remodeling and lengthening of G1 [^6,83^]. Moreover, differentiation competence itself varies within G1, with early-G1 cells preferentially initiating mesendodermal programs and late-G1 cells showing increased competence for neuroectodermal specification [^6^].

As mentioned before, MLO marker activity varied with cell-cycle, with this variation being specific to a differentiation stage (**Fig. 3d**). In ectoderm, SFPQ and SON were highest in Early S (0.76 and 0.69, respectively) and declined to negative values by G2 (-0.46 and -0.52, respectively), while PML remained stable and positive throughout (2.11 to 0.36). This inverted in endoderm, where SFPQ remained stable and positive across G1, G2 and G0 (2.01 to 1.19), while PML and COILIN dropped sharply upon exit from G1 (PML: −0.01 to −0.90; COILIN: 0.32 to −0.67) (Fig. 3d). Early mesoderm (Meso4) and mixed-identity clusters showed uniformly low MLO activity across all cell-cycle stages, with no evidence of cell cycle dependent recovery. Pluripotent clusters maintained mostly high MLO activity (PHC1, NPM1, COILIN, PML and SON) across G1, S and G2 (effect size range: 2.16 to 0.18, respectively), largely independent of cell-cycle progression; only SFPQ was comparatively cell-cycle sensitive, declining from 0.64 to −0.03 between G1 and G2. Representative images of these phenotypic interactions between differentiation, cell-cycle and MLOs are shown in **Fig. 3e**. Together, these observations significantly extend previous work [^54,57,84–88^] indicating that MLO activity is tightly coupled to cell-cycle progression and differentiation stage, with a complex pattern of interactions for all nuclear MLOs.

### Subcellular scale phenotyping resolves MLO co-occurrences and stage-dependent organelle interactions

To resolve how individual MLOs co-occur within the nucleus, we next examined the subcellular scale latent space (z_3_). Given the orthogonality of scaleVAE and the removal of cell-cycle and differentiation information from this scale relative to the latent space at the cellular scale (**Fig. 2, Supplementary Fig**. **6** and **7**), z_3_ visualization through a UMAP showed a clear structure organized by MLO markers (**Fig. 4a** and **b**). Leiden clustering of this space resolved MLO signal into well-defined clusters, with intensity for five of the six markers concentrated in a subset of clusters (Rep 1: clusters 1 4 6 and 7; Rep 2: clusters 2, 3, 4, 5 and 7) and largely absent from the rest (**Fig. 4b, Supplementary Fig. 11**). To focus subsequent analysis on nuclear regions where MLO co-occurrence could be meaningfully assessed, we retained only the MLO-rich clusters (**Supplementary Fig. 11**), further partitioning them into finer subclusters, and computed the mean intensity of the markers of every subcellular patch (see Methods). We then grouped the resulting 124 subclusters by the cosine similarity of their MLO mean intensity feature profiles, cutting the resulting dendrogram at a similarity of 0.5, which yielded eight recurring compositional states ranging from 5 to 30 subclusters each, where most (seven out of eight) states drew subclusters from both replicates (**Supplementary Fig. 12a** and **b**). Of the subclusters, we retained 82 belonging to states 1, 2, 4, 5, 6 and 7, which were present in both biological replicates and showed clear MLO expression (excluding states 3 and 8; **Supplementary Fig. 12b-d**).

**Fig. 4.**
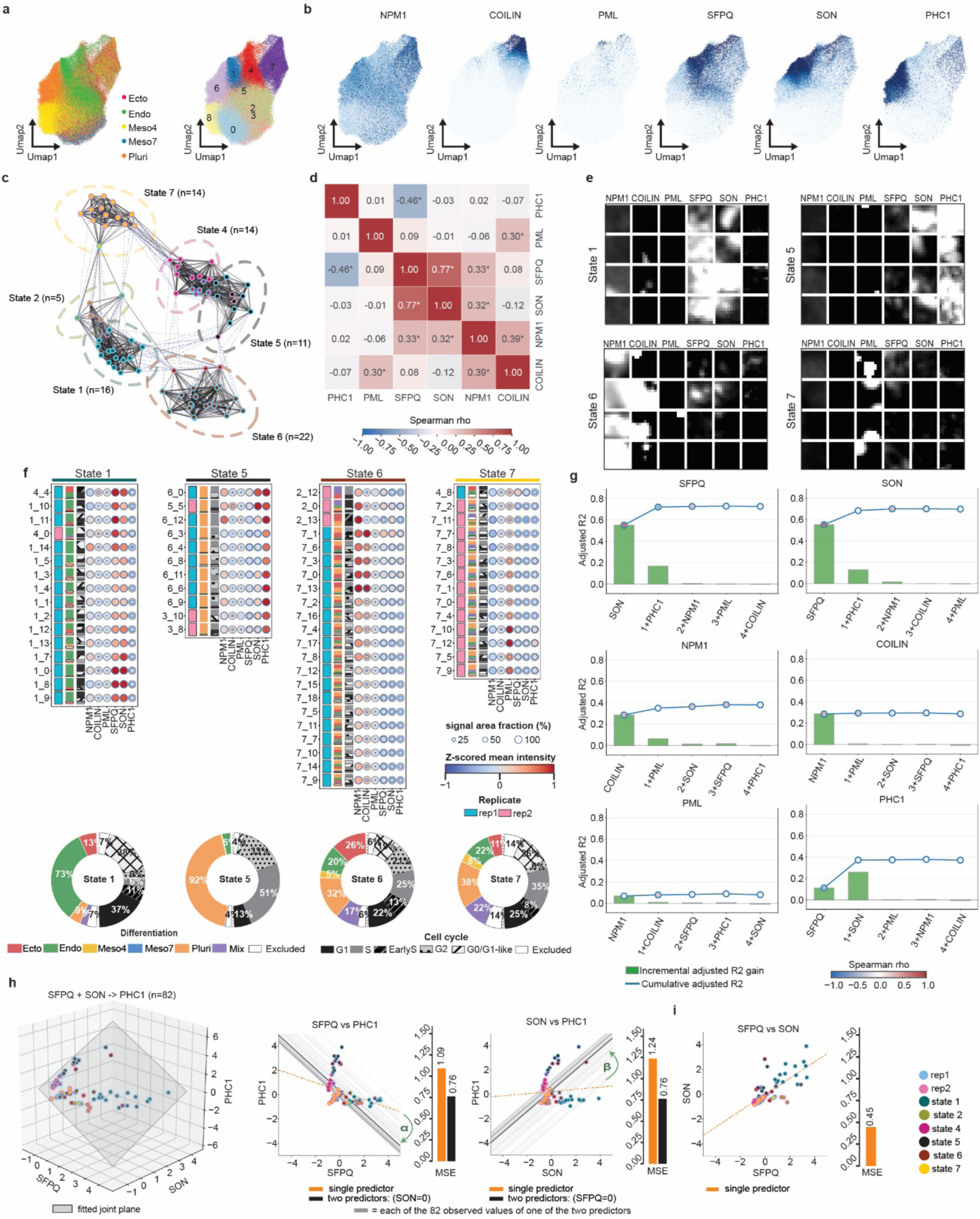
The subcellular scale latent space (z_3_) resolves MLO co-occurrence states and cross-marker predictive relationships. **a,** UMAP projections of the subcellular scale latent space z_3_ for Rep1, colored by differentiation stage (left) and by Leiden cluster assignment (right). **b,** UMAP plots of z_3_ (Rep1) colored by MLO mean marker intensity. **c,** Network representation of cosine similarities between subcellular organelle scale subclusters. Nodes are individual subclusters, colored by replicate; edges connect subclusters with cosine similarity ≥ 0.3 (dashed, 0.3–0.5; solid, ≥ 0.5), line width is proportional to similarity. Dashed ellipses indicate states obtained from this similarity structure. States 3 and 8 were excluded from the analysis (see Methods). **d,** Spearman correlation between MLO markers, computed for the subcluster’s z-scored mean marker intensity across the retained 82 subclusters retained after excluding states 3 and 8, ordered by average-linkage clustering of 1−ρ. * indicates Benjamini–Hochberg (BH) corrected p-values < 0.05. **e,** Representative subcellular scale images of MLOs for selected states (1, 5, 6 and 7). **f,** Dot plot of MLO marker intensity for each subcluster, grouped by state. Outer dot color indicates the marker z-scored mean intensity; inner dot size reflects the average signal area fraction (fraction of pixels in a patch that is above 0) of that marker within the subcluster. Side bars indicate replicate and, when available, differentiation and cell-cycle stages, colored by the patch-weighted distribution of the stages from the cellular scale clusters (Supplementary Data 4); patches whose parent cellular scale cluster was excluded from cellular scale analysis are shown as an empty portion of both bars. Donut charts below show the patch-weighted composition, pooled across all subclusters within that state. States 1, 5, 6 and 7 are shown here; states 2 and 4 are shown in Supplementary Fig. 12d, f. **g,** Incremental predictability analysis of target MLO level by remains MLO markers (see Methods and Supplementary Data 3). The plot shows the best-predictor path: starting from the single best predictor, each subsequent marker is whichever remaining predictor yields the highest cumulative adjusted R^2^ (blue line/dots) at that step, with the R^2^ gained by adding that marker shown as a green bar. Dot fill color indicates the Spearman correlation (as in d) between that added marker alone and the target. n = 82 subclusters. **h,** Three-dimensional visualization of the joint linear model PHC1 ∼ SFPQ + SON (left panel), with two-dimensional projections (middle and right) comparing the marginal regression line (orange, “single predictor”) to the joint model partial-regression line evaluated with the second marker fixed at 0 (black, “two predictors”). Gray lines show this same partial-regression relationship of the joint model evaluated at each of the 82 subclusters’ own observed values for the second marker. Mean squared error (MSE) of each model’s predictions is shown alongside. Green arrows (α, β) mark the change in slope between the single predictor (orange) and double predictor (black) regression lines for SFPQ and SON, respectively. n = 82 subclusters. This analysis is extended to two additional target markers in Supplementary Fig. 12g. **i,** Scatterplot of SFPQ vs. SON. The regression line of the linear model SON ∼ SFPQ is shown, MSE are as in h. n = 82 subclusters.

A pairwise analysis of the similarity between the remaining subclusters showed clear segregation of the identified states (**Fig. 4c**), further demonstrating their distinct phenotypic signatures. In addition, we found strong similarities of subclusters in states 1 and 2, as well as states 4 and 5, indicating that these states had related MLO signatures. For example, state 1 contains subclusters with relatively high intensity of SFPQ and SON, while in state 2 subclusters have high intensity of SFPQ only (**Fig. 4c**, **Supplementary Fig. 12e** and **f**). To investigate how MLOs correlate with one another, independent of state assignment, we computed pairwise Spearman correlations between marker intensities across all 82 retained subclusters (**Fig. 4d**). This revealed a selective pattern of positive correlation: SFPQ correlated strongly with SON (ρ = 0.77), and COILIN correlated with both NPM1 (ρ = 0.39) and PML (ρ = 0.30). PHC1 stood apart, showing no significant positive correlation with any marker and instead correlating negatively with SFPQ (ρ = −0.46) — the only significant negative pair. The specific pair wise correlation structure was lost if conditioning was not applied, showing high correlations between all MLOs (**Supplementary Fig. 12g**). We next asked whether this correlation structure was reflected in the states themselves (**Fig. 4e** and **f; Supplementary Fig. 12e** and **f**). Three states followed it directly: state 1 was high for SFPQ and SON but low for PHC1, state 6 was high for NPM1 and COILIN, and state 4 was high for PHC1 alone, consistent with PHC1 lacking a positive correlate. The remaining three states revealed interaction patterns that were missed by the global analysis. State 5 was high for both PHC1 and SON despite these two markers showing no significant pairwise global correlation (**Fig. 4d**), and states 2 and 7 isolated SFPQ and PML, respectively, as singletons despite each marker’s significant global correlation with SON and COILIN. These exceptions show that pairwise correlation alone does not fully predict which MLOs co-occur within a given state

We investigated whether these discrepancies were the result of complex multiway interactions between the different MLO markers, reasoning that pairwise correlation alone could not capture how markers jointly explain one another. To do this we fitted stepwise linear regressions, adding markers as predictors one at a time and recording the resulting gain in adjusted R^2^ for each MLO (See Methods; **Fig. 4g**). For three of the six MLO markers (NPM1, COILIN and PML), a single predictor accounted for nearly all the explainable variance by the cumulative MLO analysis (COILIN for NPM1, and NPM1 for COILIN and PML; **Fig. 4g**). For two markers (SFPQ and SON) we found simple multivariate interactions between two predictors. For SFPQ we found that SON alone explained 55.2% of its variance with PHC1 adding a further 16.9% as a second predictor. Similarly, SFPQ explained 55.2% of SON variance and PHC1 added an extra 13.1% (**Fig. 4g**). Interestingly, PHC1 showed a conditional multivariate interaction when SFPQ and SON were used as predictors. SFPQ alone explained only 11.2% of PHC1’s variance, but adding SON as a second predictor raised this to 37.2%, while SON alone explained essentially none of PHC1’s variance (adjusted R² = −0.006, **Supplementary Data 3**), indicating perdition of PHC1 intensities depends jointly on SFPQ and SON rather than on either alone.

These results suggest that the relationship between SON, SFPQ and PHC1 throughout differentiation lies on a complex hyperplane. To test this directly, we fit a joint regression plane predicting PHC1 from SFPQ and SON simultaneously across all 82 retained subclusters, visualized in three dimensions (**Fig. 4h, left**) and as two-dimensional projections comparing the multivariate fit (black/gray lines) to the univariate fit (orange line; **Fig. 4h, right**). Adding the second marker substantially improved the fit over the single-marker models; mean squared error dropped from 1.09 to 0.76 when SON was added to an SFPQ-only model, and from 1.24 to 0.76 when SFPQ was added to a SON-only model (**Fig 4h**). This reflects a suppression effect where SON’s marginal relationship with PHC1 is near zero but becomes strongly positive once the corresponding SFPQ value becomes known (β), and conversely for SFPQ (α) — hinting at non-linear relationship between the three markers driven by state 5 (92% pluripotent patches). scaleVAE allowed us to detect such interaction even though SFPQ and SON’s own pairwise correlation is strong (ρ = 0.77) and they explained each other well (mean squared error 0.45 for SFPQ predicting SON alone; **Fig. 4i**, **Supplementary Fig. 12h**). Biologically this means that in pluripotent cells nuclear speckles (marked by SON) have a strong co-occurrence with polycomb bodies given the high levels of PHC1. However, as cells differentiate, especially to endoderm, nuclear speckles tended to co-occur with paraspeckle instead (**Fig. 4f-h**).

Overall, these results show that scaleVAE can reveal organelle-to-organelle statistical interactions directly from images without the need for direct organelle detection and/or object segmentations. Furthermore, our results at the subcellular scale are consistent with the idea that MLOs do not act as isolated compartments but as an interconnected system [^89^], where spatial proximity within the nucleus creates the opportunity for functional crosstalk between them [^90^].

### scaleVAE resolves tissue domains and pathological phenotypes across disease progression in mouse kidneys

Next, we applied a two-scaled scaleVAE, consisting of a domain scale (62-by-62 pixels) and a subcellular scale (8-by-8 pixels), to analyze the pathophysiology of a immune-mediated kidney disease before and after pathological lesion formation in a mouse model [^15^] (**Fig. 5a**). These mice display a well-defined disease course progressing from acute injury to crescentic glomerulonephritis (CGN), characterized by urinary protein loss (proteinuria), followed by the formation of pathological lesions (crescents) within the renal filtering units (glomeruli), resulting in a progressive decline in kidney function [^15,91–93^]. This dataset [^15^] consists of 34 markers at a resolution of 80 nm per pixel, and 40 regions of interest (ROIs), each centered on an individual glomerulus, including controls (11 regions, 3 mice), acute phase (11 regions, 3 mice) and CGN (18 regions, 4 mice).

**Fig. 5.**
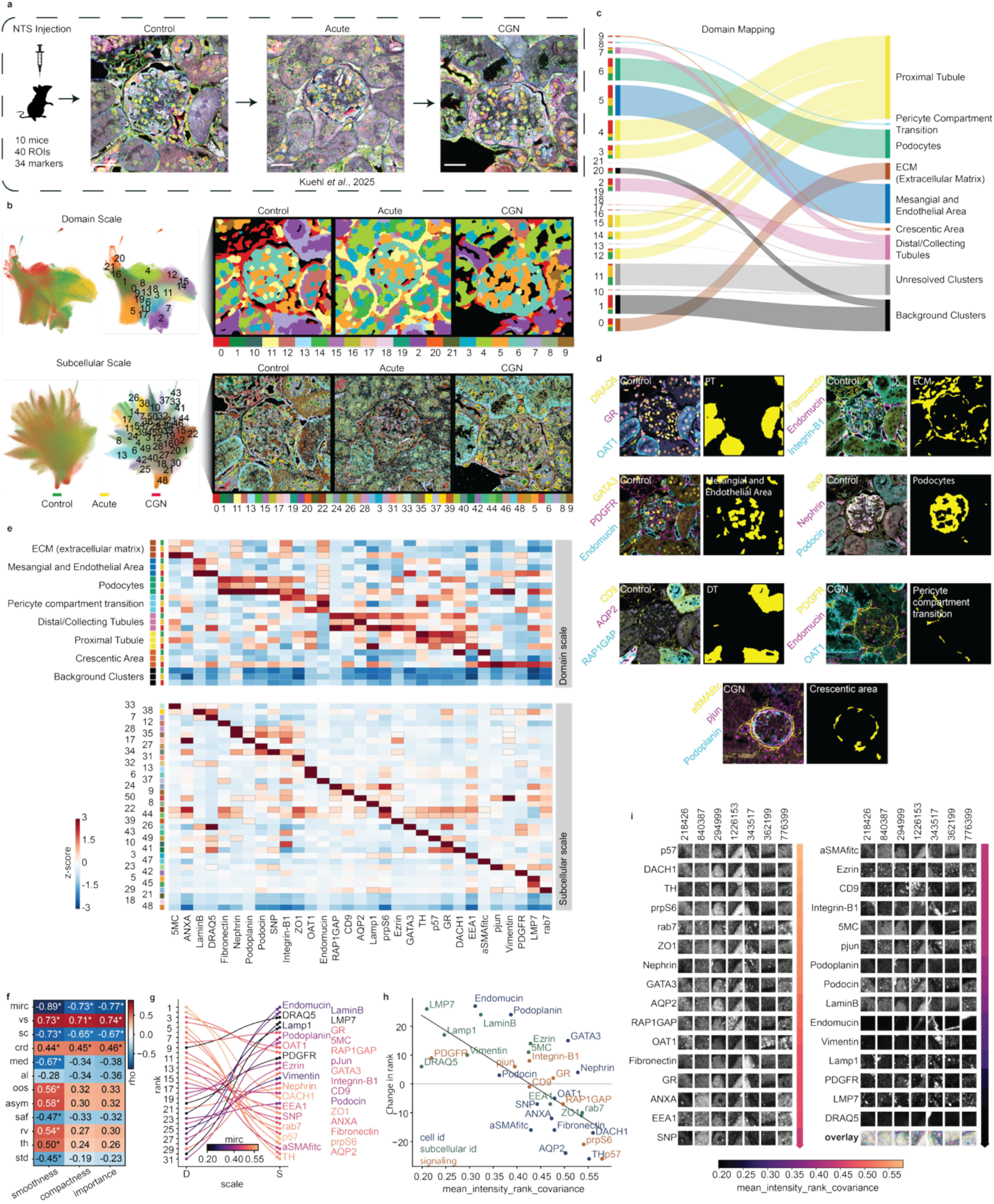
scaleVAE resolves domain and subcellular signatures in multiplexed images of tissues. **a,** Schematic overview of the experimental setup across the control, acute phase and CGN. All 34 markers included in the experiment were overlayed. Scale bar is 50 μm. **b,** UMAPs of the latent spaces of the domain and the subcellular scales together with the spatial projections of the Leiden clusters for control, acute phase and CGN examples. **c,** Sankey diagram mapping the clusters of the domain scale onto the anatomical domains of the kidney. **d,** Examples of the obtained domains across conditions with their defining markers. **e,** Conditioned stratified distribution of z-scored mean intensities of the markers across domains (upper panel). Distributions of the z-scored mean intensities of the markers across Leiden clusters of the subcellular scale (lower panel). 31 high-quality stains are shown (see Methods). **f,** Sperman correlation of the 12 computed metrics (see Methods) against the rank changes from the domain to the subcellular scale for smoothness, compactness and importance values. The color indicates the spearman’s rho. * indicates BH corrected p-values < 0.05. **g,** Ranks of markers based on their importance value at the domain and subcellular scales. The color indicates the mean intensity rank covariance (MIRC). **h,** Scatter plot of the change in rank from domain to subcellular scale against the MIRC score. Each dot is a marker; dot color indicates marker category. **i,** Example patches from the domain scale with the individual markers ordered by descending MIRC values. 7 patches shown.

Clustering the largest ‘domain’ scale latent space and projecting the clusters IDs into the pixel space (see Methods, **Suppementary Fig. 13a-d**), resulted in multiple spatially organized regions across different conditions (**Fig. 5b**). The distribution of markers (**Suppementary Fig. 13e**) and the spatial projections of clusters (**Supplementary Fig. 13f**) revealed the anatomical domains of the kidney (**Fig. 5c** and **d**) with marker specificity (**Fig. 5e**), including Podocin, Nephrin and SNP together in the Podocyte Area domain, and RAP1GAP and AQP2 in the domain of Distal/Collecting Tubules (**Fig. 5e**). At the subcellular scale, we found mainly a single dominant marker per cluster even though scaleVAE does not explicitly employ a disentanglement mechanism across markers (**Fig. 5e**).

Motivated by these observations, we asked what defines the importance of biological markers driving the domain and subcellular scale phenotypic landscapes in this dataset. To answer this, we first grouped markers into specific broad biological categories indicating cellular ID, subcellular ID and signaling pathways – mirroring the framework used in the original study [^15^], however the changes in rank of the importances between the domain and subcellular scales did not correlate with these categories (**Supplementary Fig. 13f**). We therefore computed the spearman correlation of the changes in rank of smoothness, compactness and importance to a set of 12 metrics (see Methods) to identify the axis of variation that drives the change in focus of scaleVAE (**Fig. 5f**). We found that the mean intensity rank covariance (MIRC) explains the change in ranks strongly and significantly (ρ = -0.77, **Fig. 5f-i, Supplementary Fig. 13h** and **i**). MIRC is a feature that quantifies the redundancy of a marker within the full panel by comparing the ordering of patches by a given marker intensity against all other markers. Therefore, this analysis clearly revealed that scaleVAE placed more relative importance in the domain scale to the globally trending markers whose intensity varied smoothly over pixel space, whereas non-redundant markers spatially organized in small regions dominated the latent space of the subcellular scale (**Fig. 5i**). As a consequence, we observed a hierarchical relationship between clusters at the domain and subcellular scales (**Supplementary Fig. 14**). For example, domain-scale clustering identified a single domain (Mesangial/Endothelial Area, D_C5) containing both nuclear and cytoplasmic signals and expressing mesangial and endothelial markers (Endomucin, GATA3, DRAQ5, PDGFR), whereas subcellular-scale clustering of the same tissue region resolved the nuclei within D_C5 into distinct mesangial and endothelial populations (S_C26 and S_C7, respectively) (**Supplementary Fig. 14a,e**). This could be because the domain scale patch size is 4.96 µm (62 pixels), while mesangial and endothelial nuclei can be as close as 0.72 µm, therefore their differences are resolved easier at the subcellular scale (0.64 µm, 8 pixels). Similarly, the subcellular scale clustering also revealed the structure formed by healthy parietal epithelial cells (PECs) around the glomerulus (**Supplementary Fig. 14 b,e)**, which can comprise a very narrow structure (estimated range: 0.48 µm – 1.92 µm), which is better resolved by the small patches of the subcellular scale.

To understand whether scaleVAE can reveal relevant pathology phenotypes, we carefully looked at the changes observed in the glomerular region comparing controls vs CGN (**Fig. 6**). Cluster D_C17, which is part of the Cresentic Area domain showing a spatial localization around the Glomerular Area (D_C6), is enriched in the CGN condition while missing in healthy controls (**Fig 6a-b, Supplementary Fig. 14d**). The cluster shows markedly higher pJUN (**Fig 6a-b**), consistent with the previous conclusion [] that pJUN pathway activation marks PEC transition within crescents, and is further characterized by increased signals PDPN and SNP, and low signal of Nephrin (**Fig 6b-d, Supplementary Fig. 14e**). Given its specific marker and spatial signature in CGN, D_C17 could suggest differentiating PECs acquiring podocyte-associated markers (Diff_PECs_1). Additionally, within the Crescentic Area domain we found second cluster (D_C9) likely marking PECs differentiating towards myofibroblasts (Diff_PECs_2) due to its spatial localization and high aSMA and PDGFR signals (**Fig. 6a-d, Supplementary Fig. 14d**). Overall, we observed some subcellular markers such as LMP7 and Vimentin increased with pathological progression (**Fig. 6b** and **d**).

**Fig. 6.**
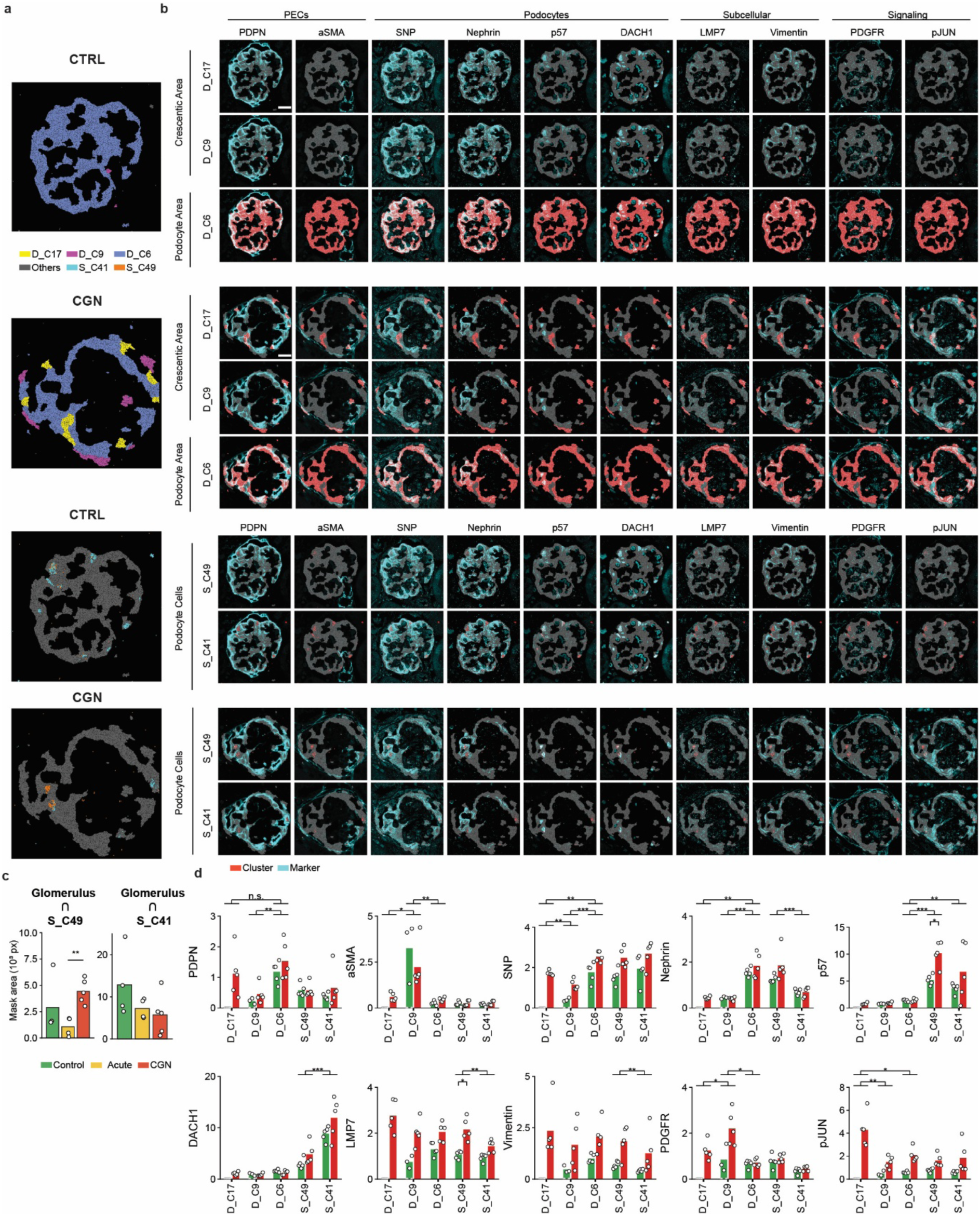
scaleVAE resolves crescentic and podocyte compartments of the glomerulus and their marker signatures in CGN. **a**, Cluster maps of one representative Control and one CGN glomerulus. Top pair, scale-0 area clusters belonging to the Crescent Area D_C17 (Diff_PECs_1, yellow) and D_C9 (Diff_PECs_2, magenta), and D_C6 (Podocyte Area, blue), with all other patches in grey. Bottom pair, scale-1 cell-level clusters S_C49 (Stressed Podocytes, orange) and S_C41 (Podocytes, cyan) on the same two crops. **b**, Each cluster mask (red) overlaid on one marker channel (cyan); white indicates pixels where the two coincide, grey the glomerulus body (union of D_C17, D_C9 and D_C6). Rows are grouped into Crescentic Area and Podocyte Area for scale 0, and Podocyte Cells for scale 1, with Control above CGN in each block. Columns group the markers as PECs, Podocytes, Subcellular and Signaling. Scale bar, 200 px. (16 µm) **c**, Mask overlap between the glomerulus and S_C49, and between the glomerulus and S_C41 (Voronoi-map pixels assigned to both masks), given as area in 10³ px. Bars, group mean; dots, individual ROIs, one per kidney slide. Samples; n = 4 Control, n = 4 Acute and n = 5 CGN ROIs. Exact two-sided permutation test on the difference of means; ** p < 0.01. No pairwise difference reached p < 0.05 for S_C41. **d**, Mean marker intensity within each cluster, computed per glomerulus and size-factor corrected. Bars, mean across glomeruli; circles, individual glomeruli. Green Control, red CGN. Small grey circles on the baseline mark a cluster absent from that glomerulus. Samples; n = 5 Control glomeruli from four slides and n = 5 CGN glomeruli from five slides. Brackets with narrow ends compare Control with CGN within one cluster; brackets with wide feet compare two clusters with both conditions pooled. Only selected comparisons are shown. *** q < 0.001, ** q < 0.01, * q < 0.05, n.s. not significant; see Methods. Each panel is scaled to its own marker.

Within the glomerulus (**Fig. 6a-b, Supplementary Fig. 14e**) the domain-scale clustering identified that Podocyte Area (D_C6) encompassed both nucleus and cytoplasmic signals characterized by high SNP and DACH1, while the subcellular scale clustering of the same region additionally resolved the nuclear phenotypes of podocytes in two distinct states; Podocytes (S_C41) and Stressed Podocytes (S_C49) (**Fig. 6a-c, Supplementary Fig. 14e**). The Stressed Podocytes cluster is characterized by high p57, high LMP7 and relatively low DACH1 compared to Podocytes (S_C41) and lacked the increase of the methylation marker 5MC observed in CGN for the Podocyte (S_C41) cluster (**Supplementary Fig. 14e**).

Although our analysis of the pathology of CGN focused on glomerular crescent formation, we were able to detect secondary biological alterations. For example, in the CGN condition we detected a domain marking a Pericyte compartment transition (D_C8) away from proximal tubules (**Supplementary Fig. 14c,e)**, and as expected we found elevated phospho-S6, marking mTOR-1 pathway activation in the tubular areas, including distal and collecting duct (D_C2, D_C7), as well as proximal tubules (**Supplementary Fig. 14e**).

Therefore, the two-scale design of scaleVAE for the mouse kidney PathoPlex dataset illustrates how the complementary resolution between domain and subcellular scales reveals the specific progression of the CGN condition. Interestingly, due to scaleVAE’s ability to extract biological information in a hierarchical manner, we observed that the subcellular scale clustering could resolve the changing signatures of signaling pathways and cellular states and help distinguish cell types at the domain scale groups, adding biological detail beyond what tissue/domain level analysis alone provides.

## Discussion

Multiplexed imaging and other spatially resolved methods allow us to quantify increasingly larger numbers of biological markers simultaneously. Therefore, these datasets are highly complex, showing specific signatures at different biological scales, ranging from the organelle and subcellular scales to the cell population and tissue scales. However, only a handful of methods try to mine such signatures, but they lack a framework to quantify and a mechanism to ensure the extractions of scale specific orthogonal phenotypes [^45–51,77^]. Here we introduced scaleVAE that automatically discovers and separates scale specific dominant sources of biological variation, enabling the learning of orthogonal phenotypic manifolds across biological scales, thus maximizing the biological information obtained from the data in an unsupervised fashion.

We benchmarked scaleVAE with a new multiplex imaging dataset of nuclear MLOs in hiPSCs and compared to existing state-of-the-art approaches (Vit-MAE and SwinViT)[^51^], which in contrast to scaleVAE, allow multiscale modeling through an additive mechanism namely attention. scaleVAE outperformed both models showing that its subtractive modeling via conditioning was better at orthogonal pattern extraction because it focused on signatures that were missing at the previous scale. Importantly, we achieved this without the need to segment cells or MLOs, nor needing to engineer specific features to measure segmented objects.

In its current implementation, the main conceptual limitation of scaleVAE is that information is passed only top-down, from larger scale to smaller scales, this limits the ability of scaleVAE to specifically and automatically select important markers at the larger scale. Also, from a technical point of view scaleVAE is not channel invariant, meaning that it requires architecture modifications and *de novo* training with each new dataset. However, owing to its compact architecture, scaleVAE can be trained rapidly on datasets of moderate scale. Future implementation of orthogonality on architectures such as vision transformers [^45,50^] should address these limitations.

The analysis with scaleVAE enabled us to perform the first systematic analysis of the changes in nuclear MLOs phenotypes throughout hiPSCs differentiation. At the multicellular scale we showed that population density itself correlates with the cell cycle and MLO levels. It is also known that spatial location of a cell within its population contributes to cell-fate decisions[^3,7^]. This latter point is likely one reason mesoderm cells showed reduced activity throughout our analyses. We lack this spatial context in our experimental model as we differentiated each germ layer separately. Physiologically, mesoderm specification depends on instructive signals from neighboring germ layers rather than occurring autonomously [^94^]. Missing these signals could reduce both the growth rate of the mesoderm lineage — resulting in a sparser population — and the general levels of MLOs due to low cellular activity.

The cellular scale provided us with the information that MLO activity is jointly governed by cell-cycle phase and lineage identity. While all MLO marker expression peaks in G1, individual MLOs diverge along germ layer trajectories as the cell cycle progresses. This observation is interesting because G1, in particular, has an impact on cell-fate decisions. Pluripotent cells have a shortened G1 phase [^83^] Endoderm specification begins in early G1, while neuroectoderm differentiation is initiated toward the end of G1[^5,6,95^]. Together with our observations, this supports a functional role for MLOs in cell-fate decisions. Previous studies have linked individual MLOs to differentiation or cell-cycle progression in isolation. For example, paraspeckles increase upon exit from pluripotency [^54,84^], nuclear speckle organization is coupled to cell-cycle progression [^57^], PML bodies peak in G1 and decline through S/G2 [^86^], and PHC1 activity is high in pluripotent cells [^88^]. With scaleVAE, we constructed a more systematic, germ-layer-resolved view of these phenotypes, showing that MLO dynamics are not uniform but highly vary in both differentiation and cell-cycle.

Our analysis in subcellular scale suggest specific inter-organelle relationships in nucleus rather than independent behavior, where spatial proximity within the nucleus could create the opportunity for functional crosstalk between them [^90^]. Whether these associations reflect causal functional dependencies remains to be tested with perturbation approaches.

Next, we tested the versatility of scaleVAE by moving from cellular-level to tissue-level data, using a mouse kidney dataset produced with PathoPlex [^15^]. As in the iPSC dataset, each scale’s latent space captured a distinct biological compartment, indicating that this scale-wise separation generalizes beyond a single tissue type. In the kidney data, the change in marker importance between the domain and the subcellular scales followed the specific spatial distribution of the markers, with markers associated with the broader regions defining the domain scale. As a result, both domain and subcellular scales provided distinct biological insights into CGN progression. This suggests that the ability of scaleVAE to disentangle scale relevant information can yield complementary, non-redundant biological insights increasing the information obtained for the data.

In conclusion, scaleVAE allows to model multiplexed imaging data with minimal human intervention, extracting scale-specific biological signatures that can be used to study the hierarchical structure of biological systems, derive new biological insights and study temporal-spatial progression of disease conditions maximizing the extracted disease phenotypes in more translational settings.

## Methods

### Culture and differentiation of human induced pluripotent stem cells

We cultured the human induced pluripotent stem cell line HMGU1 (Human Pluripotent Stem Cell Registry accession code ISFi001-A) in StemMACS iPS-Brew XF medium (Miltenyi Biotec) on six-well plates coated with Matrigel (catalog no. 354263, Corning), diluted 1:100 in DMEM/F-12 (Thermo Fisher Scientific). We passaged the cells using StemMACS Passaging Solution (Miltenyi Biotec) and seeded them in iPS-Brew XF medium supplemented with 10 µM Y-27632 (StemMACS). Starting on the following day, we replaced the medium daily with iPS-Brew XF without supplements. We passaged the cells at approximately 80% confluence.

We differentiated pluripotent cells at passages below 30 into endoderm, mesoderm, and ectoderm using the StemMACS Trilineage Differentiation Kit (Miltenyi Biotec) according to the manufacturer’s instructions. We dissociated the cells into single cells using TrypLE Express (Gibco) and seeded them onto ibidi 96-well plates (catalog no. 89626) or ibidi 24-well plates (catalog no. 82426), both coated with Matrigel (catalog no. 354263, Corning), diluted 1:100 in DMEM/F-12 (Thermo Fisher Scientific).

Cells were seeded at equivalent densities across plate formats: approximately 18,000 cells/cm² for ectoderm and mesoderm differentiation, ∼23,000 cells/cm² for endoderm differentiation, and ∼14,000 cells/cm² for pluripotent controls. This corresponded to 10,000, 13,000, and 8,000 cells per well in 96-well plates, and 27,500, 35,750, and 18,900 cells per well in 24-well plates for ectoderm/mesoderm, endoderm, and pluripotent conditions, respectively. Cells were then exposed for 7 days to the appropriate lineage-specific differentiation media specified in the kit. In parallel, we initiated early mesoderm differentiation by seeding cells at the same density on day 4 using only Mesoderm 1 medium from the kit.

### EdU labeling and fixation

At the end of day 7, we treated pluripotent and differentiated cells with EdU for 40 min using the Click-iT™ EdU Imaging Kit (Thermo Fisher Scientific). Following EdU treatment, we washed the cells with Dulbecco’s PBS, fixed them with 8% paraformaldehyde for 10 min, and stored them in PBS at 4 °C until imaging.

### Iterative indirect immunofluorescence imaging (“4i”)

We performed iterative indirect immunofluorescence imaging as previously described [^13,14^], with the following modifications and using the primary and secondary antibodies listed in Supplementary Data 5. We prepared the sBB buffer at final concentrations of 5% BSA, 364 mM maleimide, and 0.3% Triton X-100 and we eluted the sample four times with elution buffer between each 4i cycle. We diluted primary antibodies in phosphate-buffered saline (PBS) containing 1% BSA and 0.3% Triton X-100, and secondary antibodies in PBS containing 2% BSA at a 1:300 dilution. We tested all antibodies for compatibility with the elution buffer both before and after elution. For pre-elution testing, we used all primary antibodies at a 1:200 dilution and imaged them using a Zeiss Celldiscoverer 7 microscope (**Supplemantary Fig1** and **2**).

We carried out sequential incubations with single antibodies in cycles, as shown in **Supplementary Data 5**. We washed samples once with PBS between each primary antibody incubation. We initiated EdU labeling after the Triton X-100 permeabilization step and proceeded according to the manufacturer’s instructions. We detected EdU using the commercial antibody provided with the Click-iT™ EdU Imaging Kit (Thermo Fisher Scientific). We performed immunostaining for the nuclear speckle marker SON only after EdU labeling.

### Image acquisition

We used an automated Zeiss Celldiscoverer 7 microscope for pre-elution testing of antibodies. We used diode lasers at 405 nm and 488 nm with a 20× air objective for germ layer marker imaging and a 50×/1.2 NA water-immersion objective for MLO imaging. For germ layer marker imaging, we acquired z-stacks with 12 focal planes and a step size of 1 µm. For MLO imaging, we acquired images with a pixel size of 0.055 µm in the x–y plane and z-stacks with a step size of 0.3 µm and 30 focal planes. We kept acquisition settings constant across conditions.

We acquired 4i images using an automated spinning-disk confocal high-content imaging system (Opera Phenix Plus, Revvity) controlled by Harmony software (Revvity), using a 63×/1.15 NA water objective. Fluorescence was excitated at 405 nm (DAPI), 488 nm (Alexa 488), 561 nm (PhenoVue Fluor 594), and 640 nm (Alexa 647), with simultaneous acquisition of the 405/640 nm and 488/561 nm channel pairs. Images were recorded at a pixel size of 0.188 µm in the x–y plane with an exposure time of 100 ms per channel. Z-stacks comprised 30 focal planes with a step size of 0.5 µm. All acquisition settings were kept constant across experimental conditions.

### Pre-processing of images

#### Illumination correction

We leverage the statistical properties of the images acquired per channel to estimate pixel-wise illumination and gain biases [^96^]. Certain frames might have exceptionally high pixel outliers due to the technical setup. We first start by filtering these frames based on median absolute deviation (MAD) of 99,5^th^ percentile of every frame. We approximate the distribution of MAD of frames to be comparable to the standard deviation of normally distributed data. We keep every frame that lies within 3.5-fold of the z scored of the MAD scores to approximate the illumination masks per channel.

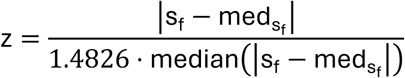

with

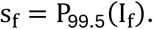

We perturb the selected frames with a gaussian noise (*σ*=50) to make them even more robust to outliers and scale them back by fitting a linear regression to the original range of pixel values. On the resulting processed frames, we compute the mean and standard deviation of intensity values at every pixel location for each channel. Illumination bias is corrected by performing per-pixel z-scoring, followed by transforming the normalized values back to the original intensity scale, yielding illumination-corrected images suitable for downstream analysis which is motivated by Stoeger, Battich *et al.,* 2015 [^96^]. The additional steps of adding gaussian noise and linearly scaling back improves upon resource requirements of Stoeger, Battich *et al.,* 2015 [^96^].

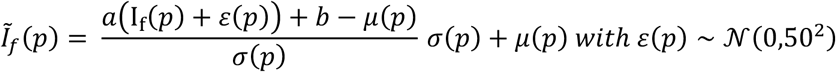

### 4i Cycle registration

Image registration is a fundamental component of correcting for systematic spatial misalignments between imaging rounds that prevent a direct comparison of markers from different cycles. We follow a 3D phase correlation approach on the range of 1^st^-99^th^ percentile of DAPI stain from each channel with a 3D Tukey window (*α* = 0.25) to reduce possible edge effects. The shift is applied to each cycle individually followed by a crop of the overlapping area to match with the first cycle’s DAPI signal (**Supplementary Fig. 4a**).

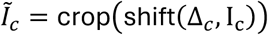

with

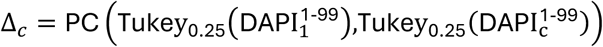

### Foreground detection

To mitigate the limitation of imperfect segmentation due to high cell density, multilayered growth, or complex tissue architecture, we reformulate the problem as a foreground detection task, treating all cellular material as foreground regardless of precise instance boundaries. For cell culture data, this approach considers cells as foreground independent of their spatial arrangement or degree of crowding, thereby removing ambiguity introduced by segmentation errors and reducing bias in downstream phenotypic analysis. We use cellpose-SAM [^97^] followed by a binarization step. For tissue imaging, we consider the whole image as foreground as there might be extracellular markers of interest (**Supplementary Fig. 4a**).

### Data cleanup

Background signal subtraction is an essential preprocessing step in microscopy imaging, as measured intensities often contain non-specific fluorescence, autofluorescence, and detector noise that are independent of the biological signal of interest. To address this, we manually estimate background signal intensities across multiple experimental conditions and subtract the measured background from all images on a per-channel basis, ensuring consistent correction across the dataset. Raw image inspection and manual background value selection were performed in ImageJ (1.54p). Following background subtraction, intensity normalization is performed to mitigate the influence of outlier pixel values that may arise from imaging artifacts or rare extreme signals by normalizing the resulting values at 99,5^th^ percentile followed by a clipping of values at 0 and 1, which stabilizes optimization and ensures numerical consistency for downstream analysis (**Supplementary Fig. 4a**).

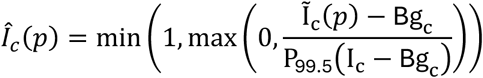

### Quality control

To remove spurious segmentations and low-quality nuclei prior to training in the MLOs dataset, we applied a data-driven intensity filter on the DAPI channel. Foreground pixels, as defined by the segmentation masks, were pooled across all images and summarized in a 256-bin intensity histogram over the normalized range [0, 1]; optionally, a uniform smoothing filter (50 × 50 px) was applied to the masked images beforehand to suppress pixel-level noise. Because this distribution is bimodal — a background peak at zero and a dominant peak corresponding to genuine nuclear signal — a threshold was set automatically at the midpoint between the two peaks, avoiding any manually tuned cutoff. Each connected component of the segmentation masks was then scored by its mean DAPI intensity, and objects falling below the threshold were removed from the masks entirely, excluding them from all downstream patch extraction and analysis. This step discards segmentation artifacts and out-of-focus or partially captured nuclei, in which the DAPI signal is not distinguishable from background (**Supplementary Fig. 4b** and **c**).

### Cell cycle analysis

#### Cell cycle classification for cVAE

For cell cycle classification used to train the cVAE model, we trained a two-scale scaleVAE model in which each scale was restricted to a distinct set of markers. The first scale incorporated transcription factor markers (PAX6, Brachyury, GATA6, and OCT4), while the second scale focused exclusively on cell cycle markers (DAPI, CDT1, KI67, and EdU). In both cases, fields of view of 80 pixels were used, reflecting the fact that cell cycle state is a single-cell–level property. The latent space learned from the cell cycle marker scale was subsequently clustered using Leiden clustering, yielding four distinct clusters corresponding to G1, S, G2/M, and non-cycling phases. Clusters were annotated based on the mean expression profiles of the cell cycle markers within each cluster. These annotations were used in the training of the cVAE as condition variables.

### Cell-cycle stage scoring

For each replicate, we z-scored the mean intensity of CDT1, EdU and KI67 across all cellular-scale clusters. Let *z*(CDT1), *z*(EdU), and *z*(KI67) denote the resulting per-cluster z-scores, and *I*(CDT1), *I*(EdU), *I*(KI67) the corresponding raw mean intensities. We computed five per-cluster scores:

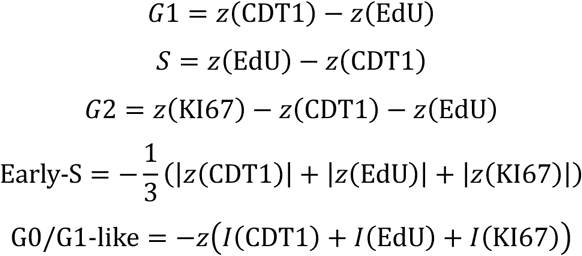

We assigned each cluster the cell-cycle stage with the highest score. Mitotic and out-of-focus clusters were excluded only when applying these assignments in downstream cellular- and subcellular-scale analyses, not from the z-scoring itself.

### Dataset generation

Prior to dataset generation, slide images were split into train and test sets, 80% and 20% respectively. The generation is based on foreground pixel sampling. For each dataset, a binary mask is used to define valid foreground pixels, which serve as candidate sampling locations. Patches are constructed by randomly sampling from this set of valid pixels, with all scales centered on the same pixel location to ensure spatial correspondence across scales. To promote diversity in the sampled content, we explicitly control the degree of overlap between the smallest-scale FOVs using a *max_overlap* parameter (0.3 for MLOs data and 0.5 for the CGN data) preventing excessive intersection while allowing limited overlap when desired based on the patch size of the finest scale, a *margin* parameter (5 for both datasets) that limits the sampling of pixels at the immediate borders of the foreground objects, *empty_patch_ratio* (0 for both datasets) parameter regulating the ratio of pixels inherent to the slide images against padded number of pixels to keep the sampling on the measured pixel space, a *fill_in_value* (0 for both datasets) for filling in the padded pixels.

### Model training

#### VAE training

We trained an unconditional variational autoencoder (VAE) on 10×10-pixel multiplexed image patches covering all 15 channels. The encoder consists of five 3×3 convolutional layers with 12, 24, 48, 96 and 192 output channels (strides 2, 2, 2, 2, 1; SELU activations, no batch normalisation), followed by two 1×1 convolutions producing the mean and log-variance of a 192-dimensional convolutional latent space; the decoder mirrors this with five transposed convolutions (192 → 96 → 48 → 24 → 12 → 15). The objective was a sigma-VAE reconstruction term, i.e. a Gaussian negative log-likelihood whose noise scale is estimated analytically from the batch residuals, plus the KL divergence to a standard normal prior with weight 1, linearly warmed up over the first 5 epochs to avoid early posterior collapse (see scaleVAE training). Models were optimised with Adam (learning rate 1×10⁻⁴), batch size 256, for 25 epochs.

#### cVAE training

We trained a conditional variational autoencoder (cVAE) on 10×10-pixel multiplexed image patches covering all 15 channels. Each patch was conditioned on the cell-level annotation of the cell it belongs to, namely cell type (5 classes) and cell-cycle phase (4 classes); both were mapped through learned embedding tables and concatenated into a single condition vector. This vector was used as the condition for the training. The encoder consists of five 3×3 convolutional layers with 12, 24, 48, 96 and 192 output channels (strides 2, 2, 2, 2, 1; SELU activations, no batch normalisation), followed by two 1×1 convolutions producing the mean and log-variance of a 192-dimensional convolutional latent space; the decoder mirrors this with five transposed convolutions (1192 → 96 → 48 → 24 → 12 → 15). The objective was a sigma-VAE reconstruction term, i.e. a Gaussian negative log-likelihood whose noise scale is estimated analytically from the batch residuals, plus the KL divergence to a standard normal prior with weight 1, linearly warmed up over the first 5 epochs to avoid early posterior collapse (see scaleVAE training). Models were optimised with Adam (learning rate 1×10⁻⁴), batch size 256, for 100 epochs.

### scaleVAE training

scaleVAE is formulated as a hierarchical generative model composed of a sequence of variational autoencoders indexed by scale *s* ∈ 1, … , *S*, where the number of scales is configurable and determined by the biological resolution of interest. At each scale s, the model defines a latent variable 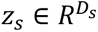 that parameterizes a continuous latent manifold and captures phenotypic variability specific to that scale. Given an input signal 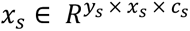 each VAE learns an approximate posterior *q_ϕ_*( *z_s_* ∣ *x_s_*, *c_s_*_–1_) and a corresponding conditional generative model *p_θ_*( *x_s_* ∣ *z_s_*, *c_s_*_–1_), where conditioning is performed on scale-specific condition variables *c_s_*_–1_ inferred from the preceding scale. At each scale, scaleVAE simultaneously learns a set of condition variables *c_s_* from the latent representation *z_s_* using a dedicated multilayer perceptron. These condition variables capture dominant axes of variation present at that scale and are used exclusively for conditioning downstream VAEs. Importantly, the latent variables *z_s_* themselves are not directly propagated across scales; instead, they serve only as inputs for learning the corresponding condition variables. This design ensures that each latent space *z_s_* encodes phenotypic variation intrinsic to its scale, while the condition variables *c_s_* account for higher-order axis of variation. Each encoder–decoder pair is implemented as a fully convolutional network, with multilayer perceptrons (MLPs) used for inference of condition variables. All VAEs are optimized jointly in an end-to-end framework, while gradient flow is restricted to within individual conditional VAE to prevent information leakage across scales. No gradients are propagated between VAEs; inter-scale communication occurs solely through the condition variables ***c_s_***. Training is initialized using a configurable head start of five epochs per scale, proceeding from the coarsest to the finest scale. Optimization is performed by minimizing, for each scale s, a variational objective consisting of a σ-VAE reconstruction term and a Kullback–Leibler divergence, with a linear warm-up applied to the KL term.

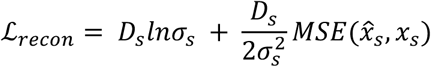

with

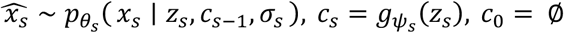

where 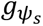 is the condition encoder of the corresponding scale. We use the analytical solution for the variance, which minimizes the (weighted) mean squared error loss (MSE) while also minimizing the logarithm of the variance [^98^]:

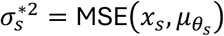

where 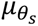 is the estimated latent mean for *x_s_*. As the prior distribution, we choose the gaussian normal distribution:

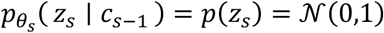

The KL term is computed as

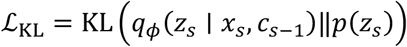

To encourage orthogonality further, we have introduced an optional term on the cross-scale correlation, which penalizes the linear dependence between the condition variable *c_i_*_–1_ and *z_i_*. Both variables are flattened, standardized feature-wise across the batch B yielding *C̃_i_*_–1_and *z̃_i_*.

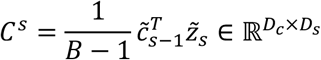

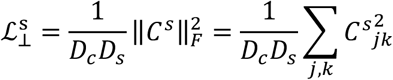

The total loss per scale is defined as the weighted sum of reconstruction and regularization terms, with equal weighting. Per scale the loss function thus becomes

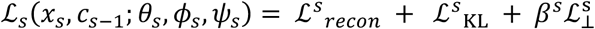

For the MLOs data, we trained a scaleVAE model, in which coarsest scale operates on 256×256-pixel patches of 2 channels (DNA and one reference channel) and consists of six 4×4 convolutional layers with 21, 28, 56, 80, 80 and 64 output channels (strides 2, 2, 2, 2, 2, 1; padding 1; SELU activations, no batch normalisation), followed by two 1×1 convolutions producing the mean and log-variance of a 64-channel 7×7 convolutional latent space; its decoder mirrors this with six transposed convolutions (64 → 80 → 80 → 56 → 28 → 21 → 2). The flattened latent of this scale (64×7×7 = 3136) is compressed by a single linear layer with SELU to a 10-dimensional condition vector, which is broadcast spatially and concatenated as extra channels to both the encoder input and the latent map of the middle scale. The middle scale operates on 80×80-pixel patches of all 15 channels (15 + 10 = 25 input channels) with five 4×4 convolutional layers of 14, 28, 56, 112 and 224 output channels (strides 2, 2, 2, 2, 1; padding 1), two 1×1 convolutions producing a 224-channel 4×4 latent space, and a mirrored decoder of five transposed convolutions (234 → 112 → 56 → 28 → 14 → 15). Its flattened latent (224×4×4 = 3584) is likewise reduced by one linear SELU layer to a 70-dimensional condition vector for the finest scale, which operates on 10×10-pixel patches of all 15 channels (15 + 70 = 85 input channels) and consists of five 3×3 convolutional layers with 12, 24, 48, 96 and 192 output channels (strides 2, 2, 2, 2, 1; SELU activations, no batch normalisation), followed by two 1×1 convolutions producing the mean and log-variance of a 192-dimensional convolutional latent space; its decoder mirrors this with five transposed convolutions (262 → 96 → 48 → 24 → 12 → 15).

For the tissue dataset, we trained a two-scaled scaleVAE model, in which the coarse scale operates on 62×62-pixel patches of all 34 channels and consists of five 4×4 convolutional layers with 64, 128, 256, 256 and 256 output channels (strides 2, 2, 2, 2, 1; padding 1; SELU activations, no batch normalisation), followed by two 1×1 convolutions producing the mean and log-variance of a 256-channel 2×2 convolutional latent space (1024 dimensions in total); its decoder mirrors this with five transposed convolutions (256 → 256 → 128 → 64 → 32 → 34). The flattened latent of this scale (256×2×2 = 1024) is compressed by a single linear layer with SELU to a 70-dimensional condition vector, which is broadcast spatially and concatenated as extra channels to both the encoder input and the latent map of the fine scale. The fine scale operates on 8×8-pixel patches of all 34 channels (34 + 70 = 104 input channels) and consists of five 3×3 convolutional layers with 64, 128, 128, 128 and 64 output channels (strides 1, 2, 2, 1, 1; padding 1; SELU activations, no batch normalisation), followed by two 1×1 convolutions producing the mean and log-variance of a 64-channel 2×2 convolutional latent space (256 dimensions in total); its decoder mirrors this with five transposed convolutions (134 → 128 → 128 → 128 → 64 → 34).

For both models trained, each scale has its own Adam optimiser (learning rate 1×10⁻⁴), both with a sigma-VAE reconstruction term and KL weight 1; the scales are switched on for training coarse-to-fine with a 5 epoch headstart for the coarser scales for 100 epochs in total.

### ViT-MAE training

The plain-ViT baseline was trained from scratch as a masked autoencoder (HuggingFace ViTMAEForPreTraining) on the same 15-channel multiplexed dataset. Patches of 260 x 260 px were sampled on the fly from the identical 3-scales with 1,000,000 patches per training epoch and 100,000 per validation epoch, margin 5 px, without any overlap and empty patches. Each patch is tokenised by a 10 x 10 non-overlapping convolutional embedding into a 26 x 26 = 676-token grid of width 512; 75% of tokens are dropped at random, the encoder sees only the visible tokens plus a CLS token, and a lightweight decoder re-inserts learned mask tokens and reconstructs the missing 10 x 10 x 15 pixel blocks under an unnormalised per-pixel MSE loss (norm_pix_loss=False). The encoder has 6 transformer layers, hidden size 512, 8 heads (64 dims/head), MLP size 2048; the decoder has 4 layers, hidden size 256, 8 heads, MLP size 1024; GELU activations and LayerNorm eps 1e-12 throughout. The model totals 23.9 M parameters. Optimisation used AdamW with learning rate 1e-3, weight decay 0.0, betas 0.9/0.999, batch size 64, gradient clipping at global norm 1.0 over 38 epochs. Latent representations were extracted from the trained ViT encoder in a single forward pass per patch. Each held-out validation patch (260 × 260 px, 15 channels; N = 238,091) was passed through the encoder in evaluation mode with the masking ratio temporarily set to zero, so that all 676 patch tokens were retained and none were replaced by mask tokens; the class token was discarded, leaving a single 26 × 26 grid of 512-dimensional tokens, each corresponding to one non-overlapping 10 × 10 px tile of the input. Since the ViT maintains a single token resolution throughout, scale-specific representations were obtained solely by spatial pooling over this one grid rather than by reading out different network depths, like the case of SwinViT. The multicellular representation was defined as the mean over all 676 tokens, integrating the full 260 × 260 px field; the cellular representation as the mean over the central 8 × 8 tokens, an 80 × 80 px region centred on the patch centre; and the subcellular representation as the single token at the grid centre, covering 10 × 10 px, hence comparable to the patch sizes of scaleVAE. For every scale, matched intensity features were computed by averaging pixels within each 10 × 10 px tiles of the input patch and applying the identical pooling operation, so that channel intensities describe exactly the same spatial footprint as the corresponding latent vector. Latents, matched channel means and patch-centre coordinates were concatenated across batches and assembled into a single AnnData object, which was treated in the same way as the scaleVAE latents in the downstream analysis.

### SwinViT training

The Swin Transformer backbone was trained from scratch in a SimMIM-style masked-image-modelling setup (HuggingFace SwinForMaskedImageModeling) directly on the 15-channel multiplexed imaging data. Training patches of 320 x 320 px were sampled on the fly from the 3-scales with 1,000,000 patches drawn per training epoch and 100,000 per validation epoch, a border margin of 5 px, without any overlap and empty patches. Each patch is tokenised by a 10 x 10 non-overlapping convolutional patch embedding (32 x 32 = 1024 tokens), and 60% of the tokens are replaced by a learned [MASK] embedding at every step; the loss is the masked-pixel reconstruction error of the decoder, a 1 x 1 convolution from the 512-dim final hidden state to 96,000 channels followed by a PixelShuffle with upscale factor 80, which reconstructs the full 320 x 320 x 15 patch exactly without padding. The encoder has four stages with depths [2, 2, 2, 2], heads [2, 4, 8, 16] (32 dims per head throughout), embedding dim 64 doubling at each of the three patch-merging steps (64 -> 128 -> 256 -> 512), window size 4, MLP ratio 4.0, GELU activations, no attention/hidden dropout, and stochastic depth increasing linearly up to 0.1 across blocks. The model totals 58.4 M trainable parameters. Optimisation used AdamW (learning rate 1e-4, weight decay 0.05, betas 0.9/0.999, constant schedule), batch size 64, gradient clipping at global norm 1.0 over 25 epochs.

The three scales for the MLOs dataset were extracted from the trained encoder in a single forward pass per patch. Each held-out validation patch (320 x 320 px, 15 channels; N = 238,091) was passed through the Swin encoder in evaluation mode without token masking. The encoder has four stages, the first three of which end in a patch merging that halves the token grid per side while doubling each token’s vector length (feature maps). This yields four token grids over the same patch: 32 x 32 tokens of dimension 64, each covering 10 x 10 px; then 16 x 16 of dimension 128 (20 px); 8 x 8 of dimension 256 (40 px); and 4 x 4 of dimension 512 (80 px). A representation at a given scale is defined by a grid level pooling pair, where pooling is either the single central token of that grid (subcellular scale) or the mean over all its tokens (cellular and multicellular scales). Three scales were used: the multicellular representation is the mean of the sixteen 80 px tokens of the final grid and therefore integrates the full 320 x 320 px field (512 dimensions); the cellular representation is the central token of that same grid, covering 80 x 80 px (512 dimensions); and the subcellular representation is the central token of the first, un-merged grid, covering 10 x 10 px (64 dimensions), hence comparable to the patch sizes of scaleVAE. Latents were taken directly from the stage outputs, prior to the encoder’s final LayerNorm, and were not otherwise rescaled or normalised. For each scale, matched raw-intensity features were computed by adaptive average pooling of the input patch to the corresponding token grid followed by the identical central-token or grid-mean operation, so that channel intensities describe exactly the same spatial footprint as the corresponding latent. Latents, matched channel means and patch centre coordinates were concatenated across batches and assembled into a single AnnData object, which was treated in the same way as the scaleVAE latents in the downstream analysis.

### Preservation of latent-space geometry across input sizes

To quantify how consistently the latent representation was organised across models applied at different input sizes, we compared the global pairwise geometry of each latent space to that of a reference model (the 10×10 model) (**Fig. 1 g**). Because all models were applied to the same cells, we drew a single random subsample of n = 2000 cells (fixed seed) and used the identical cell subset, in identical order, for every model, keeping the analysis tractable given the O(n²) number of cell pairs. For each model we computed the cosine distance between the latent embeddings of all cell pairs, *d*(*u*, *v*) = 1 − (*u* · *v*)/(‖*u*‖‖*v*‖), and vectorised the upper triangle of the resulting distance matrix 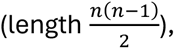 so that a given position in every model’s vector corresponds to the same pair of cells. We then quantified the similarity of each model’s latent geometry to the reference as the spearman correlation between their distance vectors, computed as the Spearman rank correlation across all cell pairs. Because it operates on ranks of pooled pairwise distances, this statistic measures monotonic agreement of the global distance structure and is invariant to the absolute distance scale; values near 1 indicate that the relative arrangement of cells is preserved across input sizes, whereas lower values indicate reorganisation of the latent geometry. We applied this comparison across scaleVAE models trained at a range of square input patch sizes (5, 6, 7, 8, 9, 10, 11, 12, 20, 30, 40, 50, 60, 70 and 80 pixels per coordinate), so that each correlation reports how the latent geometry diverges from the reference as the input size is varied. To calibrate and stress-test the metric we additionally included three control representations. As a negative control, we included the 10×10 latent space with its latent vectors randomly permuted across cells (each cell reassigned another cell’s embedding), which preserves the overall distribution of latent vectors but destroys their correspondence to cell identity; because the subsampled distance vectors are then aligned pair-for-pair against a randomly re-labelled geometry, this control is expected to yield a spearman correlation of approximately 0, confirming the metric’s lower bound of randomness. To isolate the contribution of the scaleVAE architecture from that of input size, we included a standard VAE trained at the same 10×10 input size; because the input size is held fixed, any deviation from the reference reflects architectural differences rather than resolution, namely conditioning. Finally, to probe the effect of conditioning dimensionality, we included an 80×80 model with a 1750-dimensional condition vector, allowing the change in latent geometry attributable to the condition dimensionality to be quantified independently of input size.

### Neighborhood Graph

Each scale s, the latent representations of all patches were embedded and clustered independently. The latent vectors were first reduced by principal component analysis using a randomized (Halko) solver, retaining 500 components for the multicellular, 50 for cellular, and 40 for the subcellular scales. All subsequent computations were performed in this PCA space with exact nearest-neighbor search executed on the GPU (FAISS) using rapids-singlecell library [^99^]. For each scale, 200 neighbors were considered.

### Batch-balanced neighborhood graph

Patches originating from the same slide share technical characteristics (staining intensity, section quality) that can dominate distances in latent space, a naively constructed kNN graph would connect patches primarily within slides. The exhaustive sampling of the pixel space like in the CGN data can also increase the technical effects under the same parameters of the naïve kNN graph construction. To prevent such slide-level batch effects from structuring the neighborhoods, the graph was built with the batch-balanced kNN algorithm (BBKNN; [^100^]), using the slide identity as the batch variable. Instead of selecting the k nearest neighbors globally, BBKNN connects each patch to its 8 nearest neighbors within every slide separately, so that each neighborhood spans all slides by construction. The downstream UMAP embedding and Leiden clustering therefore operate on a graph in which biological similarity, rather than slide membership, determines connectivity.

### Unsupervised clustering

Clusters were identified independently at each scale by Leiden community detection [^101^] applied directly to the k-nearest-neighbor connectivity graph of the corresponding latent space. The Leiden resolution parameter, which controls the granularity of the partition, was selected per scale by sweeping a range of values (in steps of 0.1) and inspecting the resulting partitions for stability and correspondence to known biological structure; the final analyses used a resolution of 0.8 for the multiceullar scale, 2.9 for cellular and 0.8 for the subcellular scales in the MLOs dataset. We used 1.0 and 2.2 for the domain and subcellular scales respectively in the CGN dataset. Clustering was performed with the GPU-accelerated Leiden implementation in rapids-singlecell (backed by NVIDIA RAPIDS cuGraph), with the scanpy/leidenalg CPU implementation as fallback, and a fixed random seed (42) to ensure deterministic partitions.

For the MLO data, we further performed a subclustering using the same leiden approach on a preselected set of clusters (**Supplementary Fig. 11)** based on the MLO activity to focus only on those regions of the subcellular space. The clusters were then finally subclustered individually at the leiden resolution of 1.5.

### Differential feature analysis

Upon getting the clusters on learnt latent variables per scale, we compute interpretable features on the pixel space. We then use these features to quantify the clusters and visualize the distributions in a dot plot (**Supplementary Fig. 6a**) per channel per scale. The color of the dots depicts the differences in the means of samples between the in- and out-distribution values. The larger the magnitude of the color is in absolute means, the larger the mean of the differences is. The size depicts the KL divergence between the same distributions. Both distributions were discretized innto 100 equal-width bins spanning the pooled range of the feature, normalized to unit mass, and regularized by adding a small constant (10⁻¹⁰) to each bin to avoid division by zero. The divergence was then computed as *KL*(*P*‖*Q*) = Σ_i_*p*_i_*log*(*p*_i_/*q*_i_), where *p*_i_ and *q*_i_ denote the binned in-cluster and out-of-cluster probabilities, respectively. The larger the divergence is for a feature, the more distinct this feature is for that particular cluster.

### AUC for differential cluster testing

The differential cluster analysis depends on the number of clusters, and this is inherently dependent on the interplay between the leiden resolution and the underlying latent space. To quantify the extent of this interplay, we ran the differential cluster testing across multiple resolutions for different networks. We plot the respective KL divergence and differences of means across different resolutions (**Supplementary Fig. 6b**). We then compute the area under each curve to get a global estimate of the important features across multiple leiden resolutions, accounting for different partitioning regimes. This global estimate is used to compare the sets of driving factors of individual clusters across the latent space which is depicted in percentages.

### Cluster–marker z-score heatmaps

To characterize clusters by their marker profiles, a cluster by stain matrix was built in which each entry is the mean of the per-cell mean foreground intensity of a stain over all cells assigned to the cluster. Each column of this table was then z-scored across the displayed clusters: for every channel, the mean and population standard deviation were computed over the per-cluster means themselves — so each cluster contributes exactly once regardless of its size — and the heatmap color encodes (cluster mean − mean of cluster means) / standard deviation of cluster means (**Fig. 5f** and **Supplementary Fig 13.e**). For the tissue data, Neph1, WT1 and ILR1A markers were left out from this downstream analysis following the PathoPlex paper due to poor quality. Rows and columns were arranged into an approximately block-diagonal form by ordering each cluster by the marker at which it peaks and grouping each marker under the cluster in which it scores highest, and each cluster’s defining markers were framed.

### Cluster-defining markers

To identify the markers characterizing each cluster, every stain was compared between the cells assigned to a cluster and all remaining cells. For each cluster–marker pair, we computed the mean foreground intensity inside and outside the cluster, the log2 fold change between them, *log*2((*mean_i_n* + *ε*)/(*mean_o_ut* + *ε*)) with *ε* = 10⁻⁹, and a two-sided Welch’s t-test (unequal variances) of the in-cluster versus out-of-cluster values. P values were corrected across markers within each cluster using the Benjamini–Hochberg procedure. A marker was retained as cluster-defining only if it passed both a confidence gate (corrected P < 0.05 and t-statistic ≥ 20) and a magnitude gate on the sample-size-independent effect of log2 fold change ≥ 0.5, i.e. an in-cluster mean at least 1.4 times the out-of-cluster mean. The t-statistic scales with the square root of the cluster size, so large clusters reach high confidence even for modest differences; the fold-change gate excludes such statistically confident but trivial markers. Retained markers were ranked by the product of their mean in-cluster intensity and their log2 fold change, prioritizing markers that are both abundant and specific, and all markers passing the gates were reported per cluster, so the number of defining markers adapts to each cluster. Clusters with no marker passing both gates were labeled as background clusters. In the heatmaps, a cell was framed when the marker is among the cluster’s defining markers, and its z-score in that cluster is positive (**Fig. 5f** and **Supplementary Fig. 13e**).

### Smoothness

The smoothness score quantifies how gradually a feature varies between neighboring cells in a latent space. For each scale, a k-nearest-neighbor graph was built on the latent coordinates, with edge weights decaying as *w* = *exp*(−*d*²/2*σ*²), where d is the distance between two neighbors and σ is the median of all neighbor distances in the graph. The adjacency matrix was symmetrized as *W* = ½(*W* + *W*^T^), and the unnormalized graph Laplacian *L* = *D* − *W* was formed, with D the diagonal degree matrix. Each interpretable feature was z-scored to a vector f, and its normalized Dirichlet energy was computed as 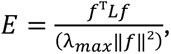 where *λ_max_* is the largest eigenvalue of *L*. This energy lies in [0, 1]: values near 0 indicate a signal that varies smoothly along the graph, values near 1 a signal that fluctuates maximally between neighbors. The smoothness score was then defined as −log₁₀(E), which is 0 for a maximally jagged feature and increases by one unit for every ten-fold decrease in energy, with no upper bound; in practice a numerical floor of 10⁻¹² inside the logarithm caps the score at 12. Higher scores therefore indicate features whose expression is organized coherently along the latent manifold rather than scattered across it (**Fig. 2e** and **Fig. 4b**).

### Compactness score

The compactness score quantifies whether the cells, in which a feature is highly expressed, occupy a single contiguous region of the latent space or are scattered across it. For each feature, the top 10% of cells by value were selected (the cells above the 0.9 quantile; ties resolved by rank, with the count n_high = ⌈0.1 · n⌉). The k-nearest-neighbor graph described above (k = 15) was then restricted to these high-valued cells, and the connected components of this subgraph were identified. The compactness score is the fraction of high-valued cells contained in the largest connected component. It is bounded in (0, 1]: a score of 1 means all high-expressing cells form one connected region of the latent manifold, while a score approaching 0 means they fragment into many mutually disconnected islands, each holding only a small share of the cells (**Fig. 2e** and **Fig. 4b**).

### Importance score

The importance score combines smoothness and compactness into a single measure of how strongly a feature is organized in a latent space and is computed as their product: *importance* = *smoot*ℎ*ness* × *compactness*. A feature receives a high importance score only if both conditions hold simultaneously — its values vary smoothly between neighboring cells in the latent space, and its high-expressing cells form a single contiguous region rather than scattered islands. Conversely, a feature that is smooth but whose high values fragment across many disconnected components, or one that is compact but noisy along the graph, is penalized multiplicatively. Since the smoothness score is unbounded above (capped at 12 by its numerical floor) and the compactness score lies in (0, 1], the importance score inherits the range of the smoothness score, [0, ∼12], and is expressed in the same units of decadic log-energy. Importance scores were computed per channel and per scale, allowing features to be ranked within each scale and their ranks compared across scales (**Fig. 2e** and **Fig. 4b**).

### Rank-change analysis

To identify which channel properties explain shifts in feature organization between scales, channels were ranked separately at each of the two scales by each of the three scores (smoothness, compactness, and importance), with rank 1 assigned to the highest-scoring channel and ties receiving the minimal rank. For every score, the rank gain of a channel was defined as its rank at a scale X minus its rank at another scale Y, so that positive values indicate a channel that rises in the ranking when the model’s field of view changes. For each channel, a panel of candidate metrics was computed from the per-cell mean foreground intensities, characterizing brightness, distribution shape, redundancy, and scale dependence. At the coarse scale, the typical brightness and spread of each channel were summarized by the median across cells (median cell intensity) as, the standard deviation across cells (intensity SD) as

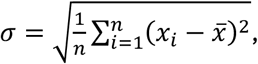

and the coefficient of variation (relative variability) as

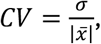

the shape of the cell-to-cell distribution was described by its skewness (asymmetry) as

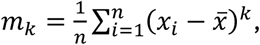

its excess kurtosis (tail heaviness) as

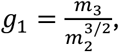

and Sarle’s bimodality coefficient as

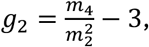

as an on/off score distinguishing channels with distinct positive and negative patch populations from gradually varying ones. The relation of a channel to the rest of the panel was captured by the mean intensity rank covariance as

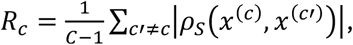

the mean absolute Spearman correlation between the channel’s ordering of cells by intensity and the orderings induced by all other channels, quantifying redundancy with the panel. Changes between the coarse and fine window were measured by three metrics: the cell ranking disagreement as

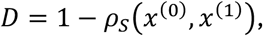

where ρ is the Spearman correlation between each cell’s value at the two scales; the signal concentration as

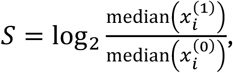

log₂ of the ratio of the fine-scale to the coarse-scale median, positive when the signal is concentrated at the patch center; and the variability shift, log₂ of the ratio of the coefficients of variation at the two scales as

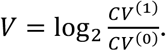

The spatial texture of each channel was quantified by its autocorrelation length: 20,000 coarse-scale patches were z-scored per channel and Hann-windowed, their two-dimensional power spectra averaged, and the radially averaged autocovariance (obtained via the Wiener–Khinchin theorem) fitted with an exponential decay exp(−lag/τ) for lags ≥ 1 pixel, with the fitted correlation length τ, in pixels, reported as the spatial scale of the stain’s texture. Finally, the signal area fraction was also computed by the fraction of foreground pixels from the mask over the total patch area. Since this constitutes a screen rather than a set of planned hypotheses, p-values were corrected across the entire grid with the Benjamini–Hochberg procedure, and for each candidate metric we additionally recorded whether the sign of its correlation was consistent across all three rank scores. Candidates were prioritized by their largest absolute correlation over the importance scores (**Fig. 5g-i**).

### Bootstrap correlation analysis of density-associated marker trends

For each replicate of the multicellular scale, we computed the observed absolute Spearman correlation, |ρ|, between per-cluster foreground fraction and each cell-cycle or MLO marker’s delta mean intensity across clusters. To estimate its sampling distribution, we resampled clusters with replacement to form a bootstrap sample of the same size and recomputed |ρ| on each resample:

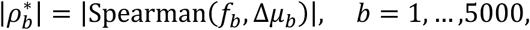

where each resample draws *n* clusters with replacement from the *n* clusters of that replicate, *f_b_* is the per-cluster foreground fraction and 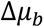 the marker’s delta mean intensity computed on resample *b*. In a complementary, non-bootstrapped analysis, we pooled both replicates and correlated per-cluster foreground fraction with the mean delta mean intensity averaged across cell-cycle markers and, separately, across MLO markers.

### Cluster-to-experimental-group assignment

For each scale, we built a contingency table of patch counts between clusters and experimental differentiation groups (Ecto, Endo, Meso7, Meso4, Pluri). From this table we computed *P*(label ∣ cluster) the fraction of a cluster’s patches originating from each group, and *P*(cluster ∣ label) the fraction of a group’s patches falling within each cluster **(Supplementary data 1)**. We assigned a cluster a pure group identity when *P*(label ∣ cluster) rounded to 1 (≥ 0.95) for a single group; we labeled clusters not reaching this threshold for any group as Mix.

### Cluster exclusion and quality-control flags

We examined example images of the patches in each cluster of cellular scale. We excluded the clusters corresponding to mitotic phase (M-phase) defined by DAPI’s characteristic condensed chromatin morphology together with KI67 positivity and out-of-focus patches were identified by zero or near-zero intensity across all markers. After cross-replicate comparisons and differential analyses we retained 31 of 38 replicate-1 and 31 of 40 replicate-2 clusters. We further flagged, but did not exclude, retained clusters as low-DAPI or as having an unclear cell-cycle assignment.

### Cross-replicate cluster similarity and matching

For each retained cellular-scale cluster, we constructed a feature vector from two quantities per marker — the delta mean intensity and the KL divergence of mean intensity — each z-scored across all retained clusters within a replicate. We computed this vector either for the full 15-marker panel (**Fig. 3b**) or restricted to the 6 MLO markers (**Supplementary Fig. 10c**). We computed cosine similarity between every replicate-1 and replicate-2 cluster pair and matched each replicate-1 cluster to its highest-similarity replicate-2 cluster (similarity ≥ 0.5). Replicate-2 clusters matched by more than one replicate-1 cluster are shown once, grouped with all matching partners (**Fig. 3c**). For the group-average comparison (**Supplementary Fig. 10b**), we first averaged cluster profiles within each combination of differentiation group and replicate, then z-scored these group-average profiles and computed cosine similarity between them.

### Effect size analysis

We defined a state as a combination of differentiation and cell-cycle stage. For each marker and each state represented among the matched cluster set (Fig. 3c; 39 clusters, comprising 25 unique replicate-1 clusters and their 14 matched replicate-2 clusters), we computed each state’s patch-weighted mean: every cluster’s z-scored delta mean intensity was weighted by its number of patches, then averaged across all clusters assigned to that state.

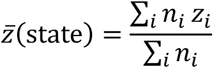

where the sum runs over all clusters *i* assigned to that state, *n_i_* is the number of patches in cluster *i*, and *z_i_* is that cluster’s z-scored delta mean intensity for the marker.

We then defined each state’s effect size as how far its patch-weighted mean departed from the average patch-weighted mean of all other states.

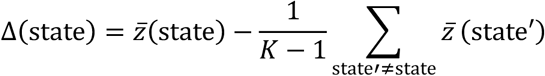

where *K* is the total number of states.

### Subcluster similarity network and state definition

We represented each subcellular-scale subcluster (n = 124) by its mean intensity for each of the 6 MLO markers, z-scored across all subclusters. We computed cosine similarity between all subcluster pairs,

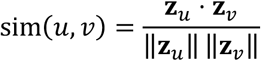

where **z***_u_* and **z***_v_* are the z-scored marker-intensity vectors of subclusters *u* and *v*, and converted it to a distance,

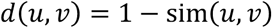

before grouping subclusters by average-linkage hierarchical clustering. Cutting this dendrogram at a distance of 0.5 defined 8 discrete states (sizes 16, 5, 12, 14, 11, 22, 14 and 30 for states 1 to 8). We excluded states 3 and 8 from correlation, incremental feature contribution and joint linear model analyses: state 3 comprised replicate-2 subclusters exclusively and did not replicate in Rep1; state 8 was a large, compositionally mixed, low-confidence state. We retained both states in the full marker dot plots and composition donuts.

### Cross-marker Spearman correlation

We computed Spearman correlations between every pair of the 6 MLO markers’ z-scored mean intensities across the selected 82 subclusters of subcellular scale retained after excluding states 3 and 8 and assessed significance by FDR (Benjamini–Hochberg) correction across all 15 marker pairs (threshold < 0.05) (Fig. 4d).

### Incremental feature contribution analysis

For each MLO marker as target, we fit all linear regression models (ordinary least squares) using 1 to 5 of the remaining MLO markers as predictors across the 82 retained subclusters of subcellular scale (31 combinations per target, 186 models in total), using z-scored mean marker intensities:

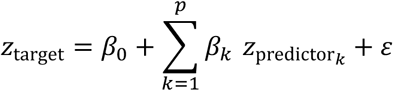

where *p* ∈ {1, … ,5} is the number of predictors in the model. Model fit was quantified by the adjusted *R*^2^,

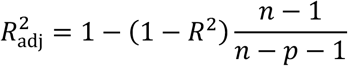

where *n*n is the number of subclusters (82) and *p*p the number of predictors. We constructed a forward stepwise path greedily, selecting at each step the predictor whose addition maximized adjusted *R*^2^. We report the adjusted *R*^2^ gain at each step,

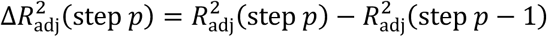

and the cumulative adjusted *R*^2^, and color each added predictor by its Spearman correlation with the target (Fig. 4g).

### Joint linear regression analysis

For a given target marker and a pair of predictor markers (e.g., PHC1 as target, SFPQ and SON as predictors), we fit a joint linear model of the target on both predictors’ z-scored mean intensities across the 82 retained subclusters of subcellular scale, defining a fitted regression plane:

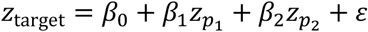

We evaluated each individual predictor’s partial-regression relationship with the target in two ways: with the second predictor fixed at 0, its z-scored mean (“with second predictor”),

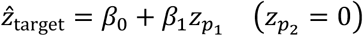

and, for each subcluster *i*i, with the second predictor fixed at that subcluster’s own observed value while *z_p_*_1_ varies continuously across its plotted range (gray lines),

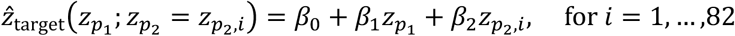

We compared these against the corresponding single-predictor (marginal) linear regression of the target on that predictor alone (“without second predictor”),

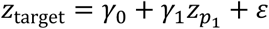

and computed mean squared error (MSE) between each model’s fitted values and the observed target values across all 82 subclusters,

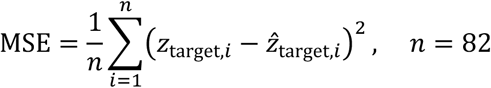

### Domain Definition

The domains for the CGN data were defined by looking at the cluster-defining markers and the spatial distribution of the cluster projected onto the pixel space. Multiple clusters were compared across different conditions to combine them into domains regardless of the condition specific patterns (**Fig. 5c**).

### Size-factor normalization

Per-slide differences in global staining brightness of the tissue data were removed before any downstream comparison. For each slide s, a single size factor sf(s) was defined as the mean, over all channels c, of that slide’s mean per-cell foreground intensity, *sf*(*s*) = *mean_c_*(*mean_i_*_∈*s*_*I_ra_*_j_(*i*, *c*)). Each cell’s per-channel mean foreground intensity was then divided by the size factor of the slide it came from, 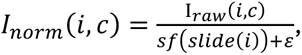, with *ε* = 1*e* − 9 guarding against division by zero. Because the same scalar is applied to all channels of a slide, relative marker profiles within a cell are preserved while the slide-level brightness component is removed.

### Identification of spatially unspecific clusters

The spatial coherence of each cluster was quantified per slide using a one-versus-rest Moran’s I. For each slide, a k-nearest-neighbor graph (k = 15) was constructed from the patch coordinates, restricting neighborhoods to individual slides. The distance-weighted adjacency matrix W was row-normalized to sum to one, so that Wv gives the neighborhood average of any patch-level quantity v. For each cluster, patch membership was encoded as *x_i_* = 1for patches assigned to the cluster and *x_i_* = 0otherwise and centered as *z_i_* = *x_i_* − *x̄*, where *x̄* is the fraction of patches on the slide belonging to the cluster. Member patches therefore carry positive values and non-member patches carry negative values in proportion to their deviation from the cluster’s expected share. Moran’s I was computed as *I* = (*n*/*S*₀) · (*z*^T^*Wz*)/(*z*^T^*z*), where n is the number of patches and S₀ the sum of all weights. The term *z*^T^*Wz*is the covariance between each patch’s membership and the neighbourhood average of its spatial neighbours, and is normalized by the variance *z*^T^*z*, yielding a correlation-like statistic approximately bounded by [−1, 1]. Values approaching 1 indicate that a cluster forms contiguous spatial domains, whereas values near 0 indicate spatially random arrangement. Per-cluster values were averaged across slides (omitting slides lacking the cluster) and are reported as mean ± s.d. Clusters with a mean Moran’s I ≤ 0.1 were classified as spatially unspecific and excluded from downstream analysis (**Supplementary Fig. 13c and d**).

### Glomerular area selection and statistical analyses

Glomerular area selection. Analysis was restricted to nine slides, four Control and five CGN; the four early-injury slides were excluded. Glomerular area selection was made on the basis of the podocyte cluster D_C6. Each analysed patch centre of that cluster was painted as a filled disc of radius 3 px, and the glomerular bounding box was taken as the 3rd to 97th percentile of the resulting extent with a 140 px margin. Isolated podocyte patches lying in tubular tissue were excluded from the box computation but retained for measurement.

Slide 2 contains three glomeruli rather than one, and was separated by two straight cuts at y = 740 and x = 1,200. The two complete glomeruli below and above the horizontal cut were retained as separate samples; the third, clipped by the frame corner, was discarded. The cuts were enforced on the measurement masks, so no patch contributes to two glomeruli. Ten glomeruli were obtained: five Control, from four slides, and five CGN, from five slides.

Intensity measurement. For each glomerulus, cluster and marker, the patch centres assigned to that cluster were painted as discs of radius 3 px and intersected with that glomerulus’s region, and the mean pixel intensity of the marker channel was taken within the resulting mask. This yields one value per glomerulus per cluster per marker. Values were corrected for per-slide brightness using the size factor described above.

Statistical analyses. The unit of replication throughout was the glomerulus, not the patch; patch centres within one glomerulus are repeated measures of a single tissue and were never treated as independent observations.

Control was compared with CGN within a cluster by two-sided Welch’s t-test on the per-glomerulus corrected means (n = 5 Control, n = 5 CGN) except for D_C9 (n = 3 Control, n = 5 CGN) with Benjamini–Hochberg correction across the 40 testable cluster–marker pairs.

Clusters were compared with one another on the pooled Control and CGN glomeruli by two-sided paired t-test on log₂ corrected intensity, paired within glomerulus. For each glomerulus the difference log_2_(cluster A) − log_2_(cluster B) *y* was formed, and the mean of these differences was tested against zero, with Benjamini–Hochberg correction across all 100 cluster pairs. The number of pairs ranged from 5 to 10 glomeruli, since each comparison uses only the glomeruli in which both clusters are present.

D_C17 was absent from every Control glomerulus, so the Control versus CGN comparison could not be performed for it; it contributes to the cluster-versus-cluster comparisons through the five glomeruli in which it is present. Because two of the five Control glomeruli derive from the same slide, the Control-versus-CGN tests were repeated with those two averaged into a single sample; no conclusion changed.

The two families of tests were corrected separately, each across its own set of comparisons. Only selected significant comparisons are annotated in the figure; the complete pairwise results are given in Supplementary Data 6. Significance is reported on the corrected value as ***q < 0.001, **q < 0.01, *q < 0.05.

Analyses were performed in Python 3.11.5 with NumPy 1.24.3, SciPy 1.11.1, pandas 2.1.4, stats models 0.14.0 and scikit-image 0.20.0.

## Contributions

A.O.C, G.M., and N.B. conceptualized the project and orthogonal phenotyping workflow. A.O.C. developed the scaleVAE. G.M. performed required experimental procedures, including full differentiation experiments and staining. A.P., G. M., F. T. and N.B. provided critical feedback during development of scaleVAE. G.M. designed and optimized data collection. G.M. and A.G.S. performed initial optimizations of 4i. G.M., K. S., I.R. and K.H. performed 4i imaging and data acquisition for the MLO datasets. A.O.C., A.P., G.M., F.T. and N.B., conceptualized the analysis of the MLO dataset. A.O.C., A.P., G.M., M.K., V. G.P., F.T. and N.B. conceptualized the analysis of the kidney dataset. A.O.C., G.M., M. K., V. G. P., and N.B. interpreted results from the analysis of the kidney dataset. A.O.C. and G.M. performed downstream analysis for all datasets. F. T., N.B., supervised analysis. A.O.C and G.M. contributed equally as co-first authors. N.B. supervised the whole project. A.O.C, G.M., and N.B. wrote the manuscript, with input from all authors.

## Code availability

The code will be made public upon acceptance.

## Data availability

Data is available upon request.

## Acknowledgements

This research was funded by the ERC Starting Grant (DynaDiff, Project # 101040548). A.O.C and A.P. are supported by the Helmholtz Association under the joint research school Munich School for Data Science (MUDS). The authors thank A. Scialdone for helpful comments on data analysis, S. Buddharaju, C. Pütz, A. Ettinger, E. Rusha, A. Feuchtinger, G. Gut and F. Pecori for comments and help on experimental set up. A.C. Rodriguez, T. Scharwz-Romond, M.E. Torres-Padilla, M. Elsner and members of the Battich and Theis labs for comments on the manuscript.

