## Supplementary Information for "Orthogonal phenotyping from multiplex imaging across scales"

This file contains Supplementary Figures 1 to 14.

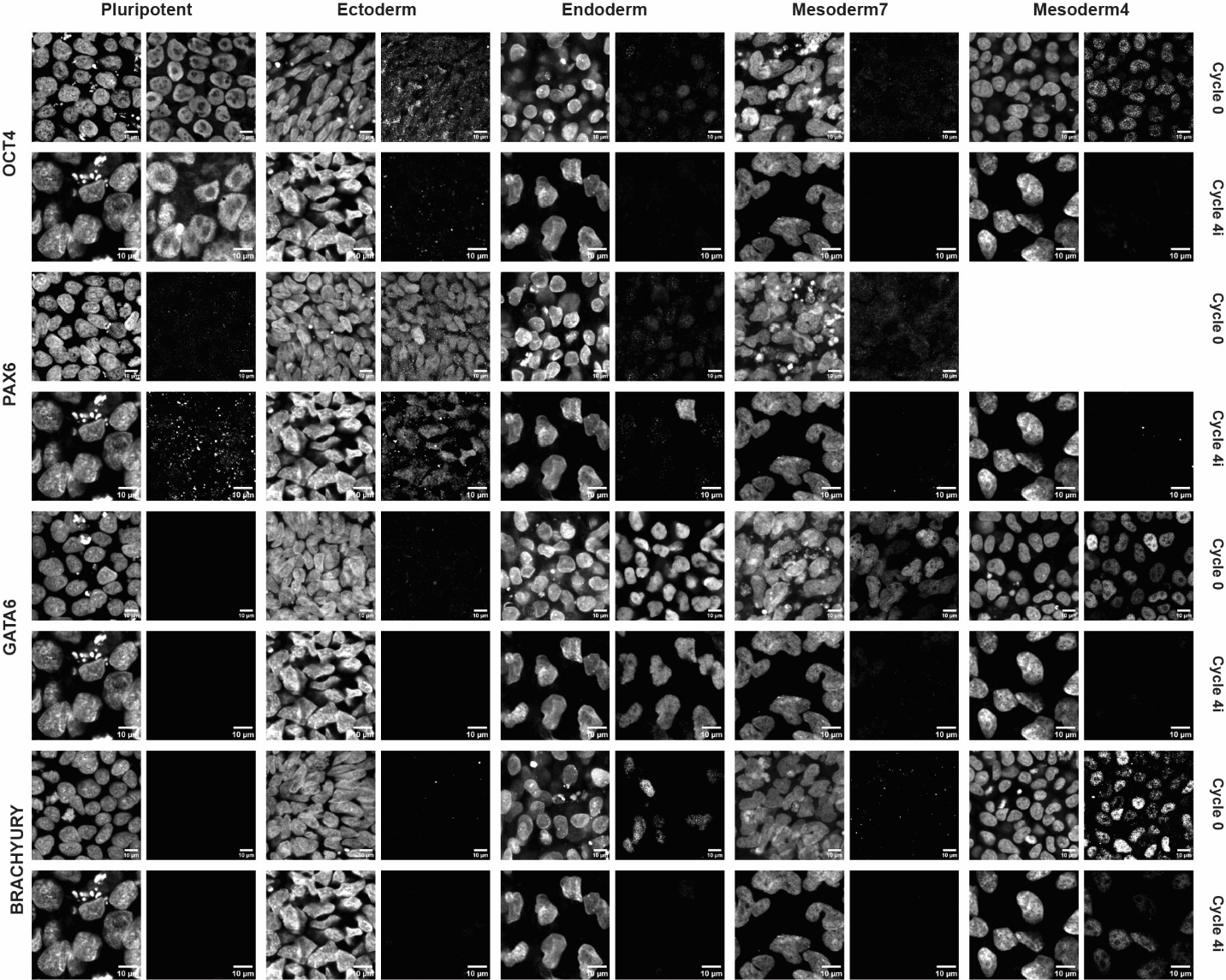

**Supplementary Fig. 1: Iterative immunofluorescence (4i) preserves marker detection across differentiation states.** Representative images comparing conventional immunostaining (cycle 0) with 4i-acquired images (cycle 4i) are shown for pluripotent and lineage-specific markers: OCT4 (pluripotent), PAX6 (ectoderm), GATA6 (endoderm), and BRACHYURY (mesoderm). Columns indicate differentiation conditions, while rows correspond to marker staining. For each marker, DAPI and antibody channels are presented as image pairs. The rightmost labels indicate imaging cycles. Scale bars: 10µm

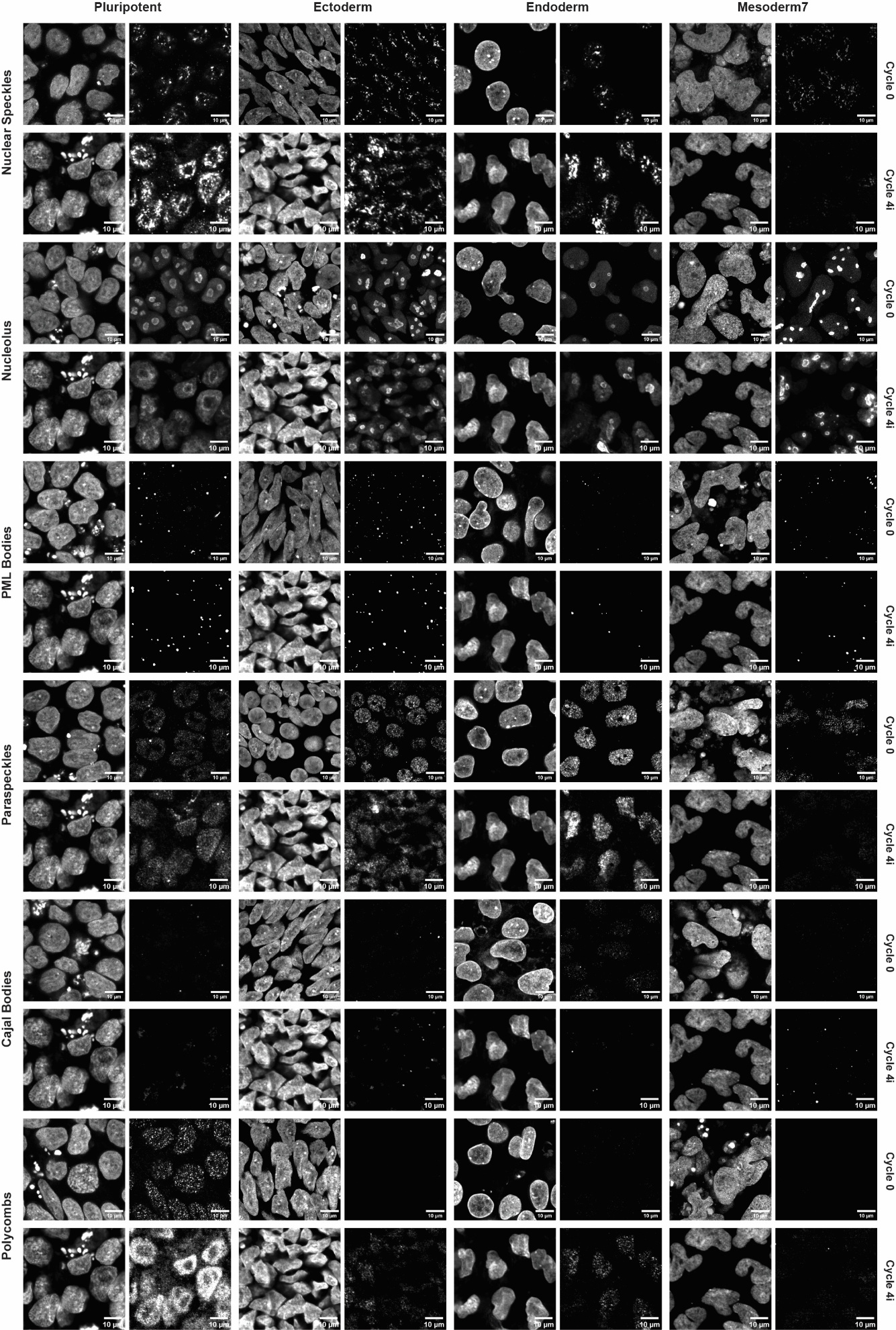

**Supplementary Fig. 2: Iterative immunofluorescence preserves detection of membraneless organelles across multiple staining cycles.** Representative images comparing cycle 0 (conventional immunostaining) and cycle 4i demonstrate consistent detection of nuclear compartments across pluripotent and differentiated states (ectoderm, endoderm, mesoderm). Rows correspond to membraneless organelles, including nuclear speckles (SRRM2, cycle 0; SON, cycle 4i), paraspeckles (SFPQ), Cajal bodies (COILIN), nucleoli (NCL, cycle 0; NPM1, cycle 4i), PML bodies (PML), and Polycomb bodies (PHC1). Columns indicate differentiation states. For each condition, DAPI and corresponding antibody staining are shown as paired images. Cycle information is indicated on the right. Scale bar: 10 µm

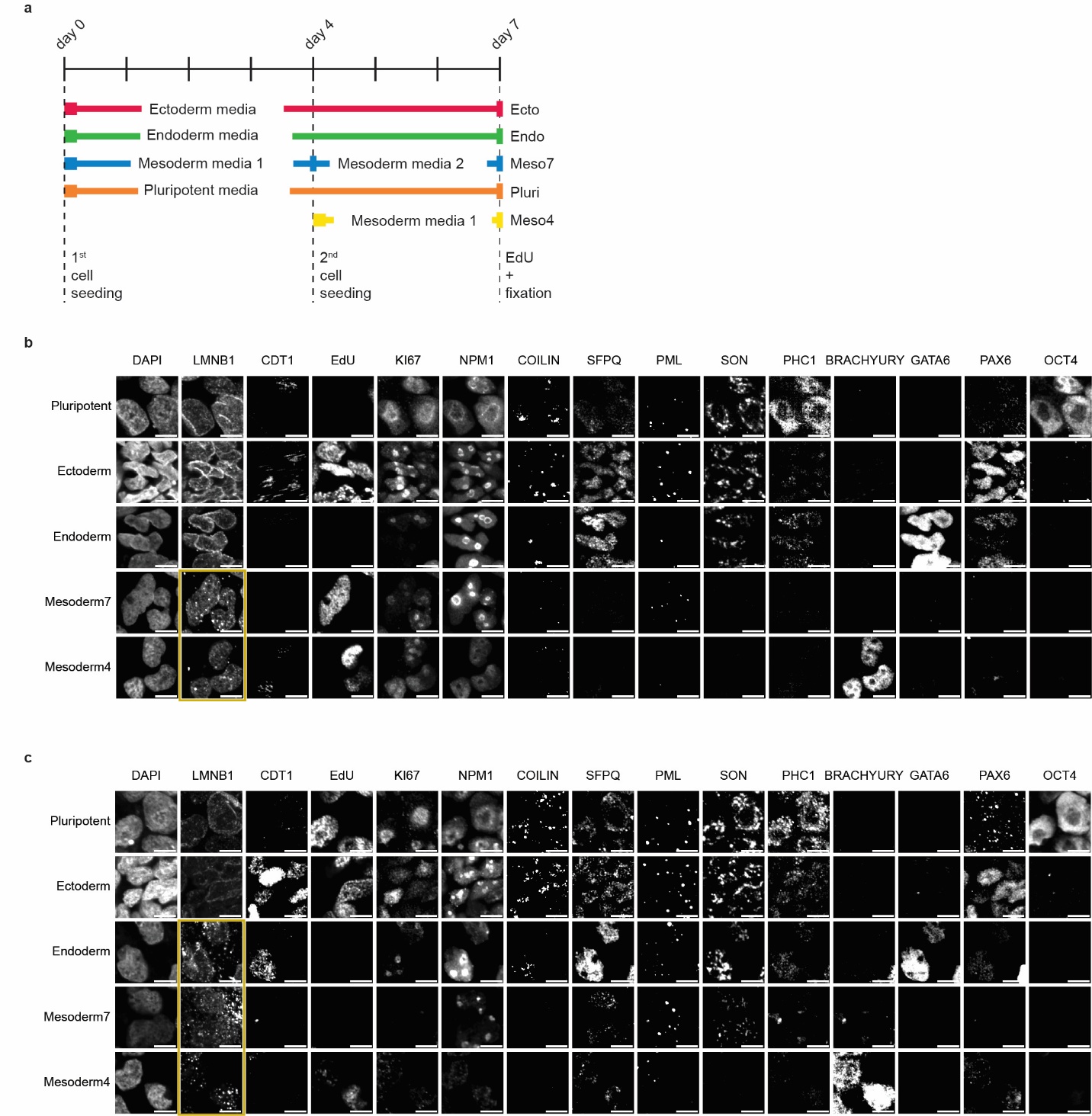

**Supplementary Fig. 3: Experimental overview.** (a) Experimental design for cellular differentiation. All conditions were established in parallel by seeding cells at day 0 in a multiwell plate. Differentiation into ectoderm, endoderm, and mesoderm (day 7) was induced by applying lineage-specific media, while pluripotent cells were maintained in standard culture media. A second mesoderm condition (day 4) was generated by seeding iPSCs at day 4. All samples were labeled with EdU at day 7 and fixed with 8% paraformaldehyde prior to imaging. (b-c) Representative images from 4i multiplexed imaging of replicate one (b) and replicated two(c) across seven cycles comprising 15 markers, including cell cycle, differentiation, nuclear structure, and membraneless organelle markers (indicated above). Rows correspond to differentiation states and columns to individual markers. For each condition, DAPI and antibody channels are shown as image pairs. Noisy signals are framed in a yellow box. Scale bar: 10 µm.

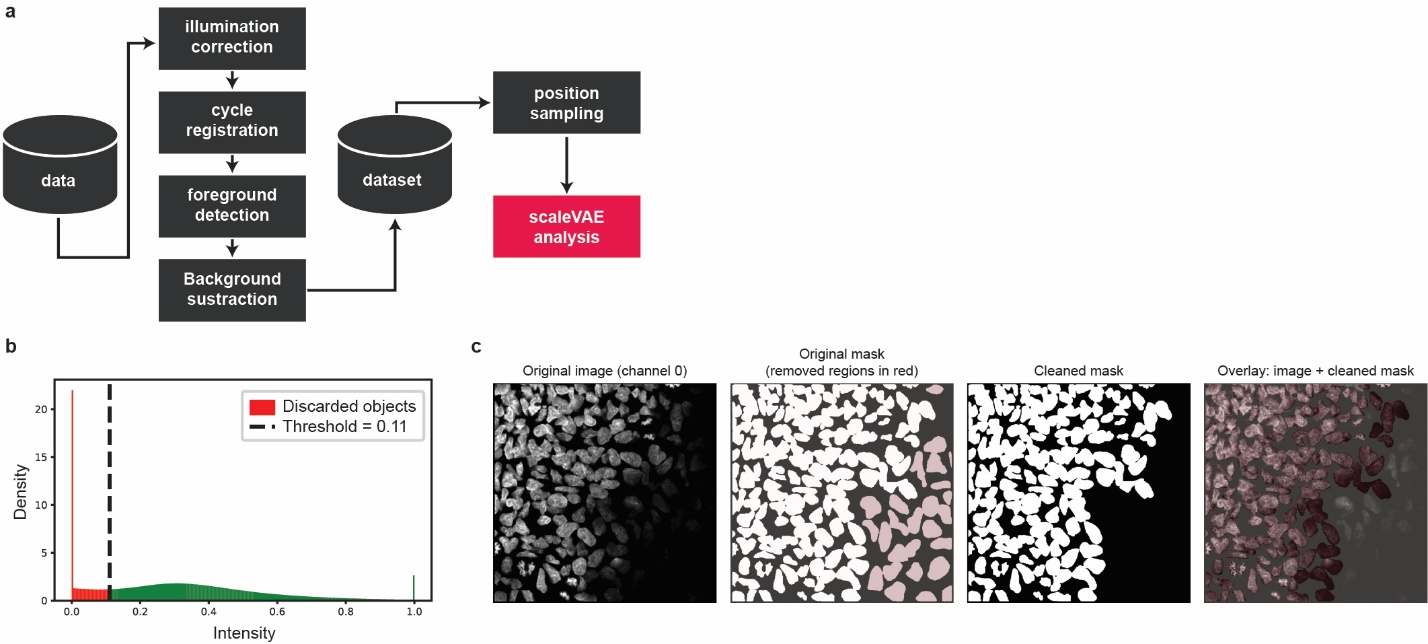

**Supplementary Fig. 4: Image preprocessing and sampling region selection.** (a) Schematic of the image preprocessing workflow applied to iterative immunofluorescence (4i) datasets, including illumination correction, cycle registration, foreground detection, and background subtraction, followed by dataset assembly, position sampling, and downstream analysis using scaleVAE. (b) Mask intensity filtering. Segmentation masks were refined by removing dim connected components whose signal was closer to background than to true fluorescence. For each DAPI (nuclear reference) image, we pooled the intensities of all foreground pixels (mask == 1) across the entire dataset and computed their histogram (256 bins over the normalized intensity range [0, 1]). Prior to computation of the histograms, images were smoothed with a uniform (box) mean filter of size 50 px to suppress pixel-level noise and stabilize peak detection; this smoothing was used only to derive the threshold and did not alter the image or mask data. A single global intensity threshold was defined as the midpoint between the background level, assumed at intensity 0, and the dominant foreground peak, taken as the center of the tallest non-zero histogram bin (threshold = peak_foreground / 2). Each mask was then decomposed into connected components (scipy.ndimage.label), and any component whose mean DAPI intensity (computed on the unsmoothed image) fell below this threshold was removed from the mask. The resulting filtered masks were used for all downstream analyses. (c) Example images on low-DAPI mask removal.

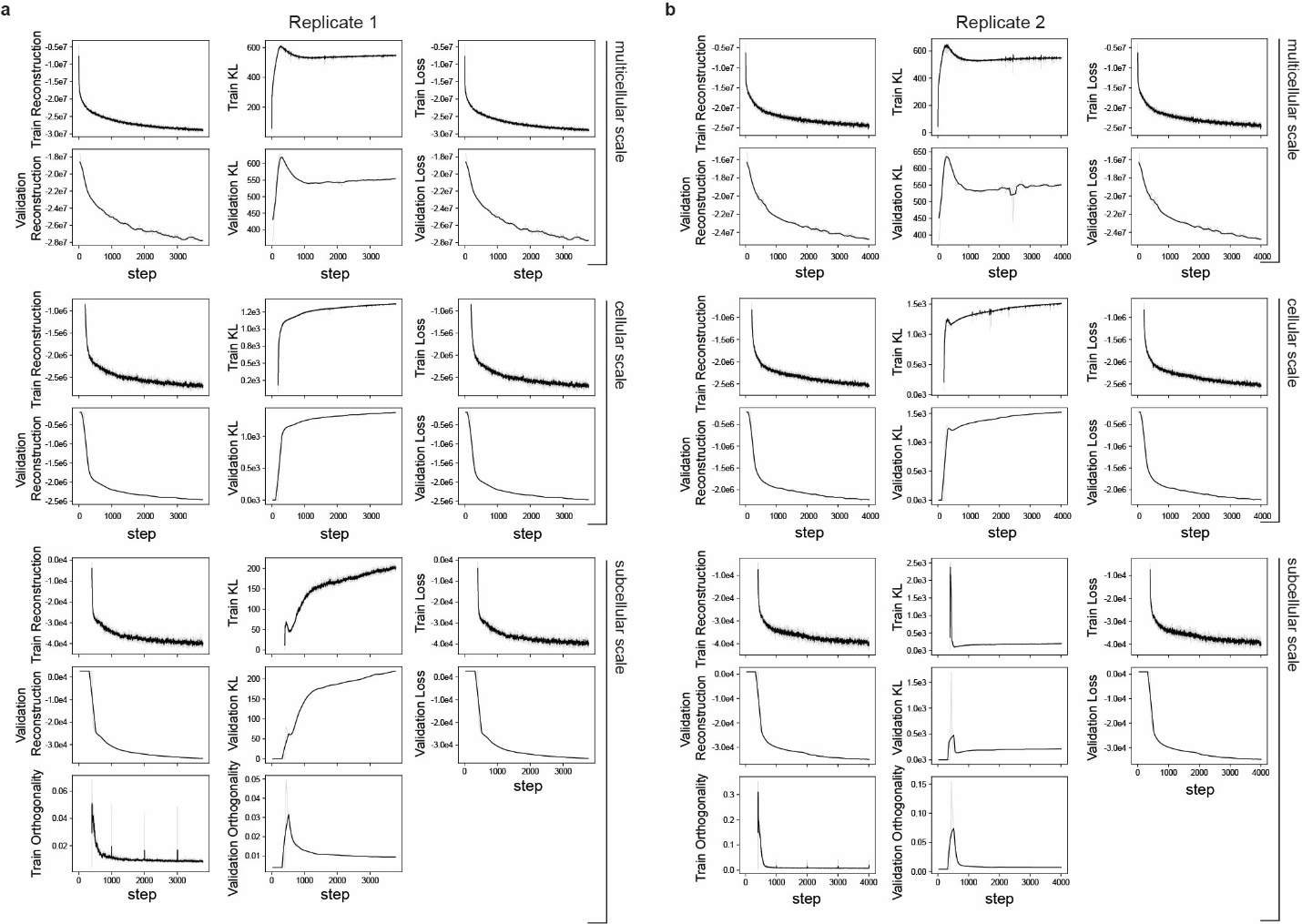

**Supplementary Fig. 5: scaleVAE training. a,**Training curves of a scaleVAE trained on the replicate 1 data. The metrics are plotted against steps of the training, i.e. total sample size over batch size. The curves are smoothened with a running window of 5 steps. **b,** Training curves of a scaleVAE trained on the replicate 2 data.

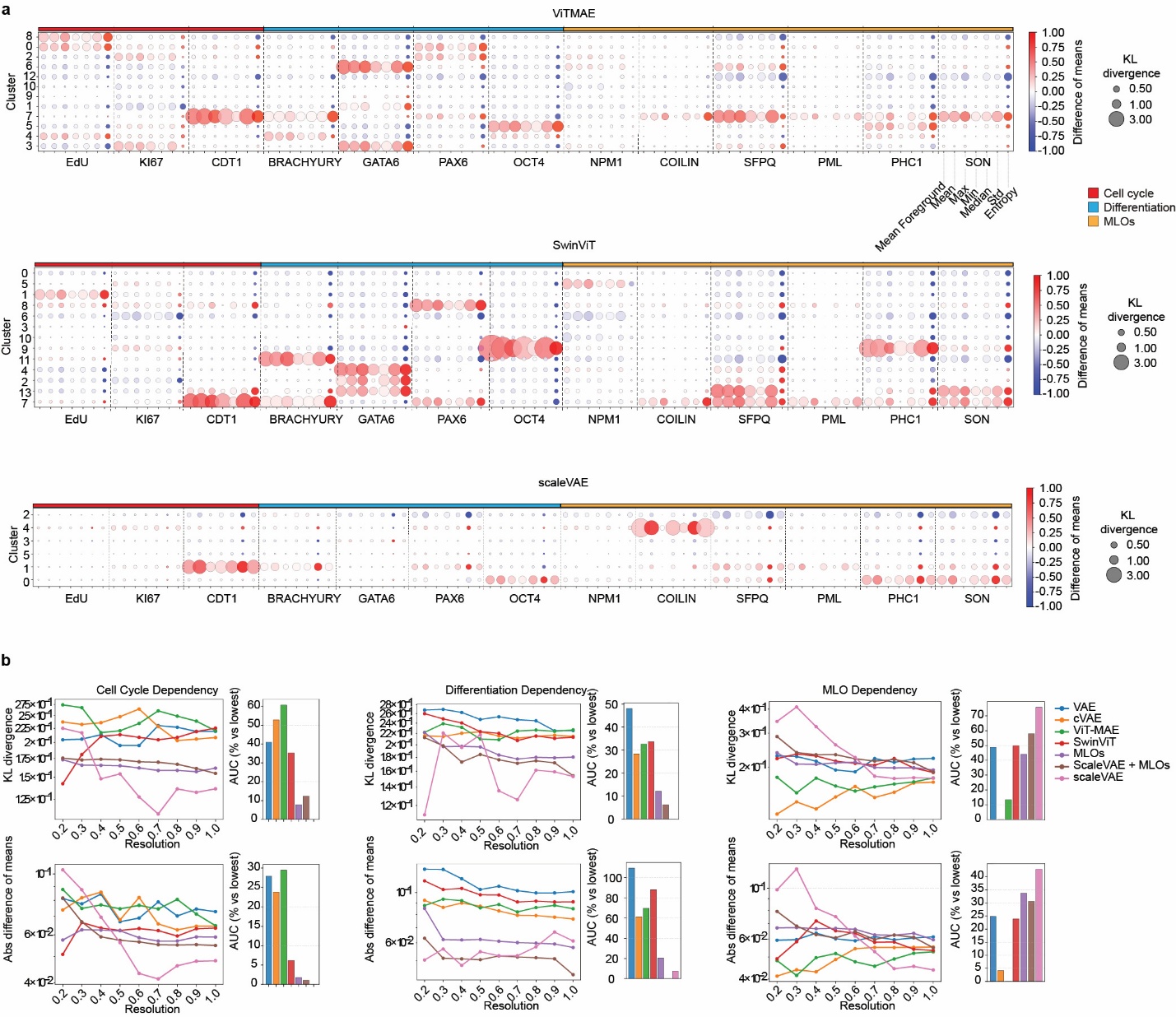

**Supplementary Fig. 6: cluster-wise orthogonality evaluation. a,** The dotplots of leiden clusters at resolution 0.7 for ViTMAE, SwinViT and scaleVAE respectively. The colors are the differences of the means and the size of the dots depitcs the KL divergence of each feature individually accordingly with Figure 2b. **b,** The first row quantifies the median of the KL divergence of the respective category of stains across different resolutions. The bar plots are the percental gain of the area under the curve normalized by the value of the smallest bar. The second row depicts the same statistics for the absolute differences of the means. The set of features are limited to cell cycle, differentiation and MLO stains from left to right columns. The lower the KL bars are, the less difference there is of that particular set of stains among the clusters across multiple resolutions of the leiden algorithm.

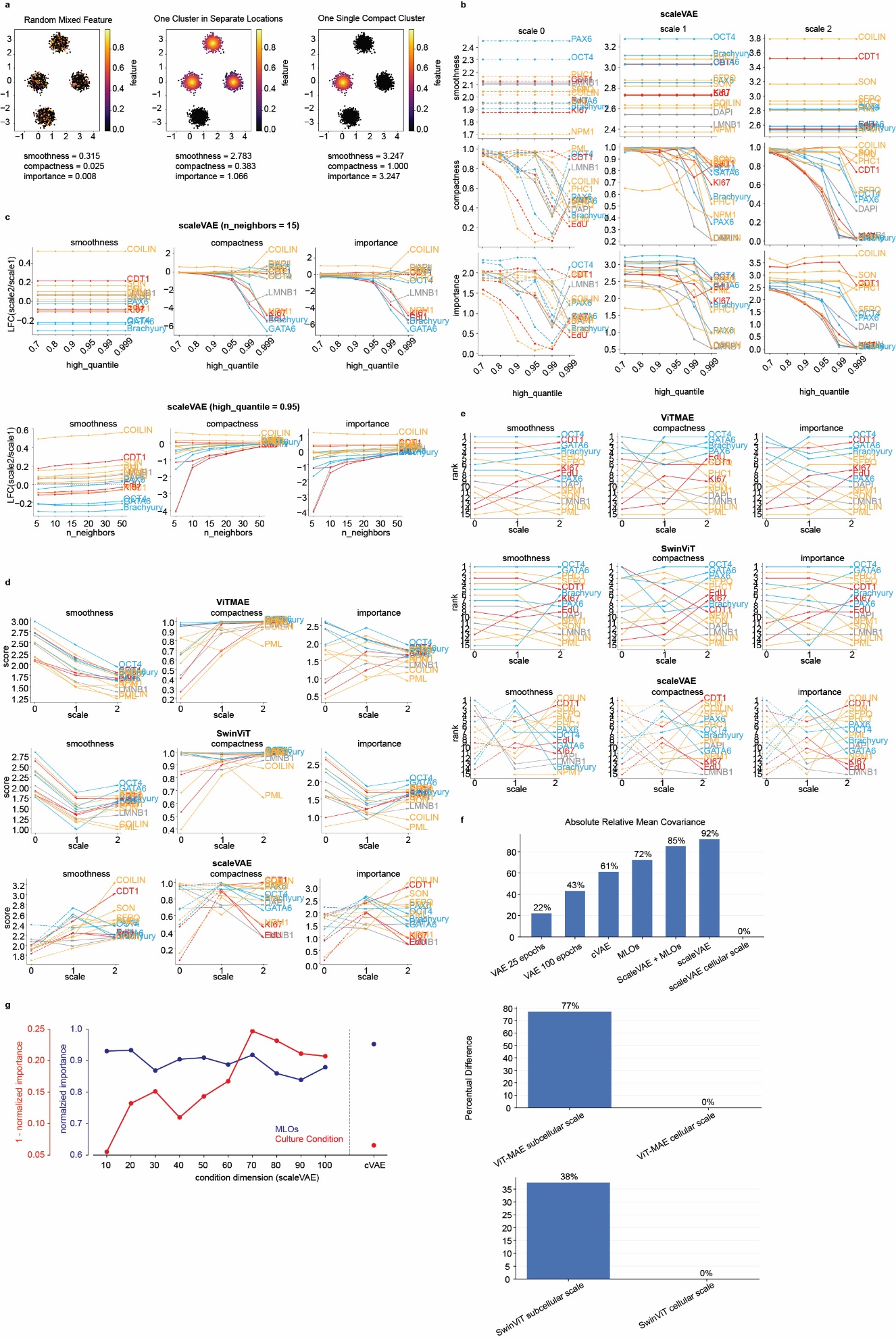

**Supplementary Fig. 7: cluster-free orthogonality evaluation. a,** Depicts the computed values of smoothness, compactness and the importance scores on a toy dataset with 3 different distributions of the same feature. For all three scores, the higher the better. **b,** Benchmarking of the high quantile value (k% in Figure 2c.) over smoothness, compactness and the importance for the 3 scales of scaleVAE. The x ticks are percentile values and correspond to 71428, 47619, 23810, 11905, 2381 and 239 samples respectively. **c,** Benchmarking the log2 fold change of the smoothness, compactness and importance scores of subcellular scales over the cellular scale at a fixed 15 neighboors for the knn graph computation. The quantile values in the x axis correspond to that of figure c. The second row fixes the quantile at 0.95 percentile and benchmarks the effect of increasing n_neighbors for the knn graph. **d,** the smoothness, compactness and importance scores at multicellular (0), cellular (1) and subcellular (2) scales for ViTMAE, SwinViT and scaleVAE. **e,** The ranking of the stains based on their computed scores in figure d. **f,** The percentual difference of orthogonality loss (see Methods) measured between subcellular and cellular scales of a VAE trained for 25 and 100 epochs, a conditional VAE conditioned on cell type and cycle, a VAE trained on MLO markers (6), a scaleVAE trained on MLO markers in the subcellular scale, a scaleVAE, a ViT-MAE and a SwinViT trained for 100 epochs on all 15 markers **g,** The normalized importance score on the average of MLO stains in blue and the complement of the importance score on the average of the transcription factor (TF) stains across different condition dimensionality of scaleVAE from 10 to 100 with n_neighbor = 15 and high_quantile = 5 at the subcellular scale. The rightmost after the dash line is a cVAE that had the culture conditions as OHE vectors that was learnt as a 1000-dimensional lookup table (nn.Embedding) during training to regress out the effect of the TF stains on the subcellular scale. For both of the lines, the higher the better.

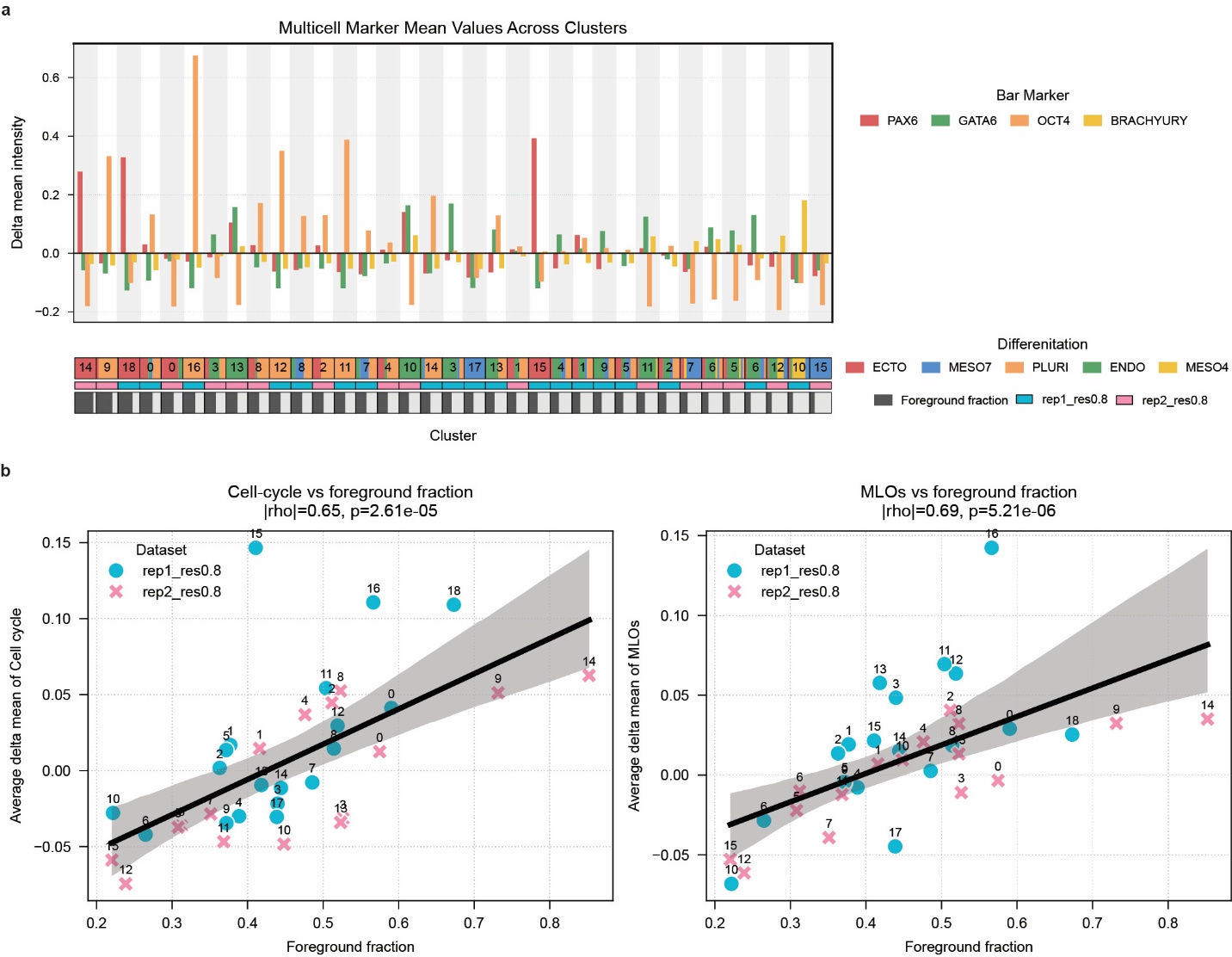

**Supplementary Fig. 8 Differentiation marker levels and density-associated trends in cell-cycle and MLO markers across multicellular-scale clusters.** **a,** Delta mean intensity of differentiation markers for each multicellular-scale cluster, computed separately within each replicate and shown combined (replicate 1 and replicate 2), ordered by descending foreground fraction, as in Extended Data Fig. 1b. Side bars indicate the germ layer/pluripotency composition, replicate, and foreground fraction of each cluster (as in Extended Data Fig. 1b). **b,** Per-cluster average delta mean intensity of cell-cycle markers (left) and MLO markers (right), plotted against foreground fraction, for all multicellular-scale clusters from both replicates combined. Line and shaded band show a linear regression fit with 95% confidence interval; |ρ| and p indicate the Spearman correlation coefficient and associated p-value (see Methods).

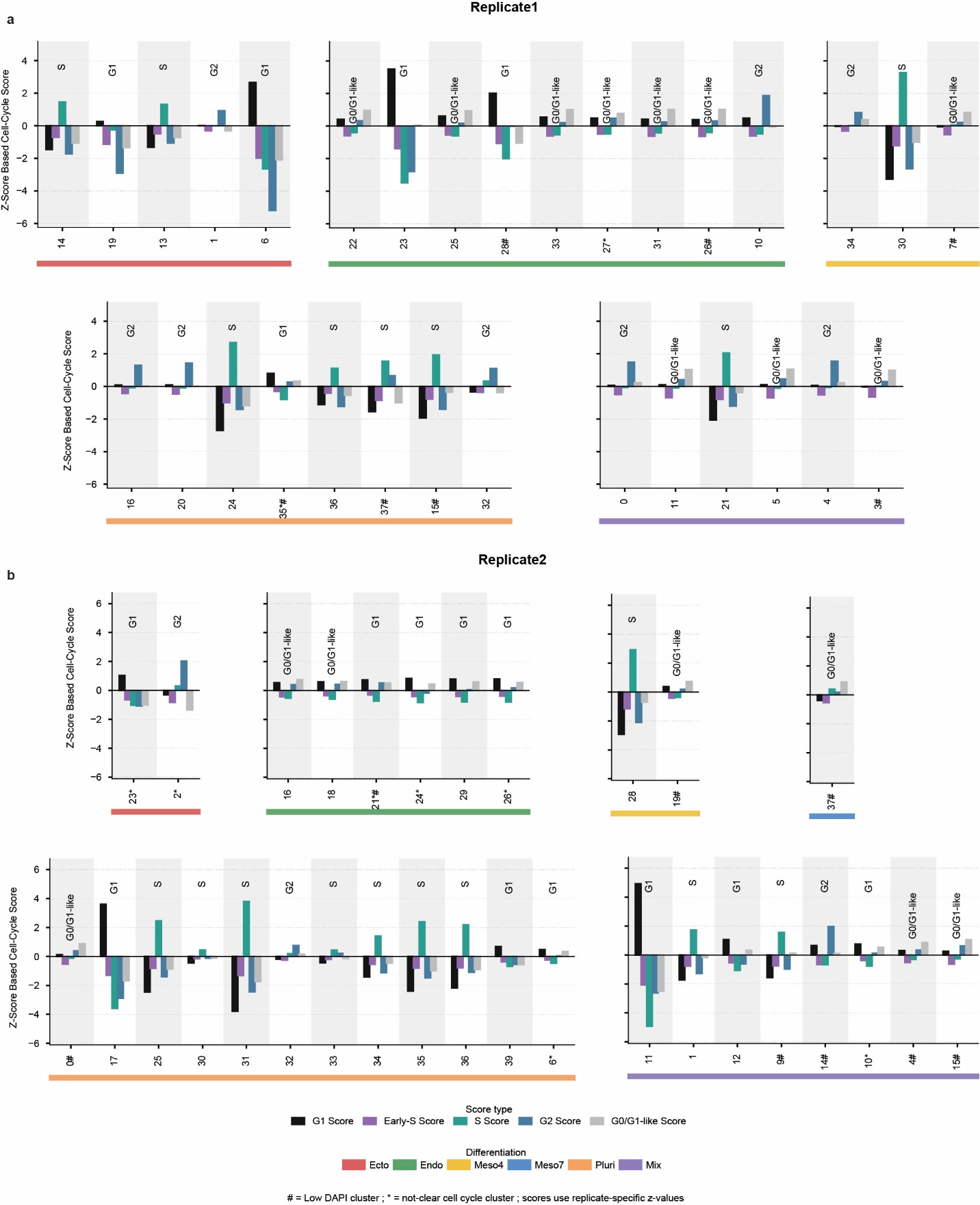

**Supplementary Fig. 9: Cell-cycle scores underlying cluster-level cell-cycle assignments in the cellular-scale analysis.** **a,** Replicate 1 and **b,** Replicate 2. For each cellular-scale cluster, five scores are shown: G1, Early-S, S, G2 and G0/G1-like. These scores are based on the z-scored mean intensity of CDT1, EdU and KI67 (see Methods). Clusters are grouped into differentiation blocks, indicated by the colored bar beneath each block. The label above each cluster's bars indicates its highest-scoring state. Mitotic and out-of-focus clusters are excluded. # denotes a low-DAPI cluster; * denotes an unclear cell-cycle assignment.

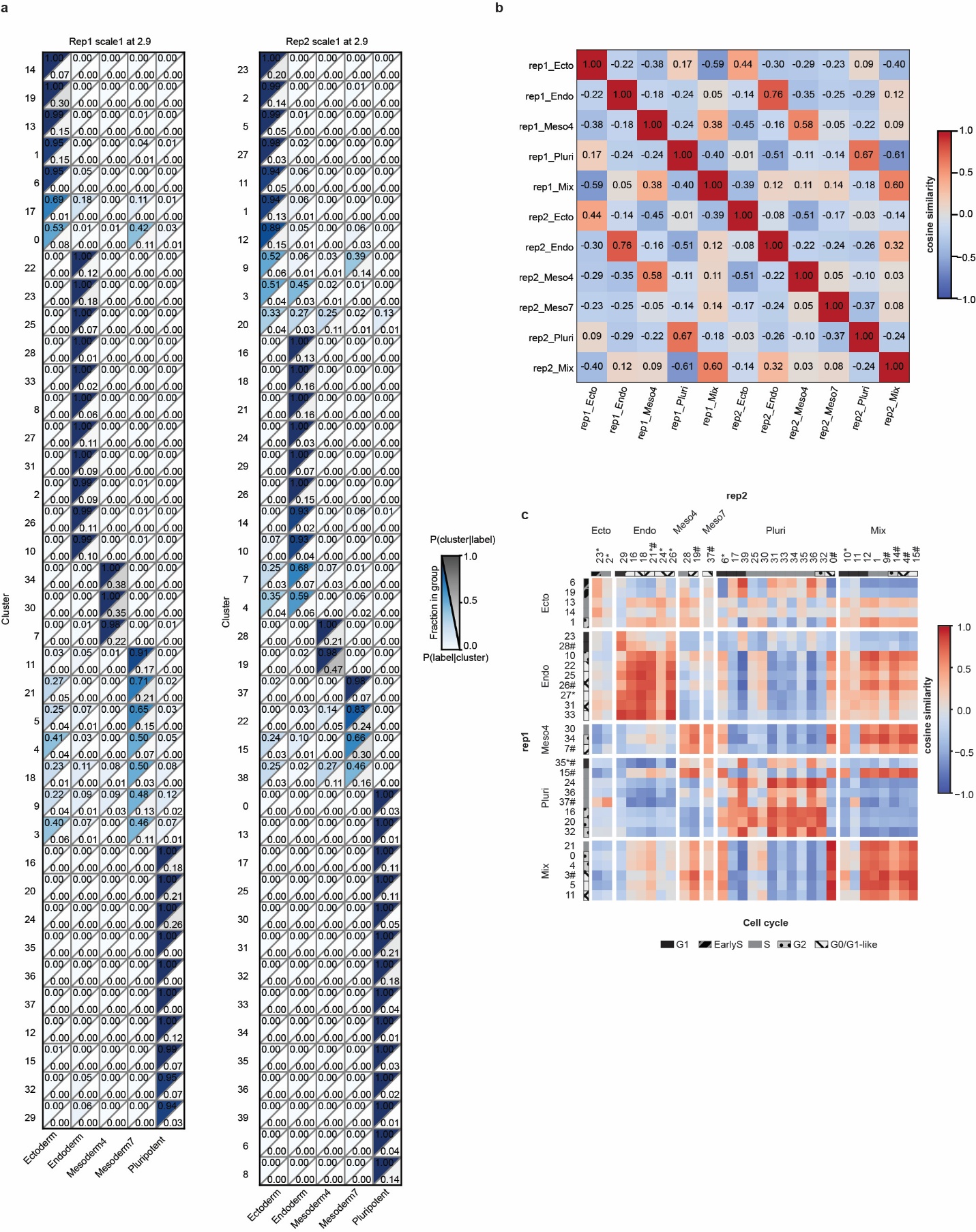

**Supplementary Fig. 10: Cluster-to-experimental-group assignment and robustness of cross-replicate cluster correspondence to marker subset.** **a,** Bipartite correspondence between cellular-scale clusters (rows) and experimental differentiation groups (columns), for replicate 1 (left) and replicate 2 (right). Each cell shows the fraction of patches in a cluster originating from a given group (P(label|cluster), blue) and the fraction of patches from a given group falling within that cluster (P(cluster|label), grey; see Methods). **b,** Cosine similarity between germ layer/pluripotency group-average profiles of replicate 1 and replicate 2. **c,** Cosine similarity between individual cellular-scale clusters of replicate 1 and replicate 2, computed using only MLO markers rather than the full marker panel (see Methods). Clusters are grouped and ordered by germ layer/pluripotency identity; icons indicate cell-cycle state.

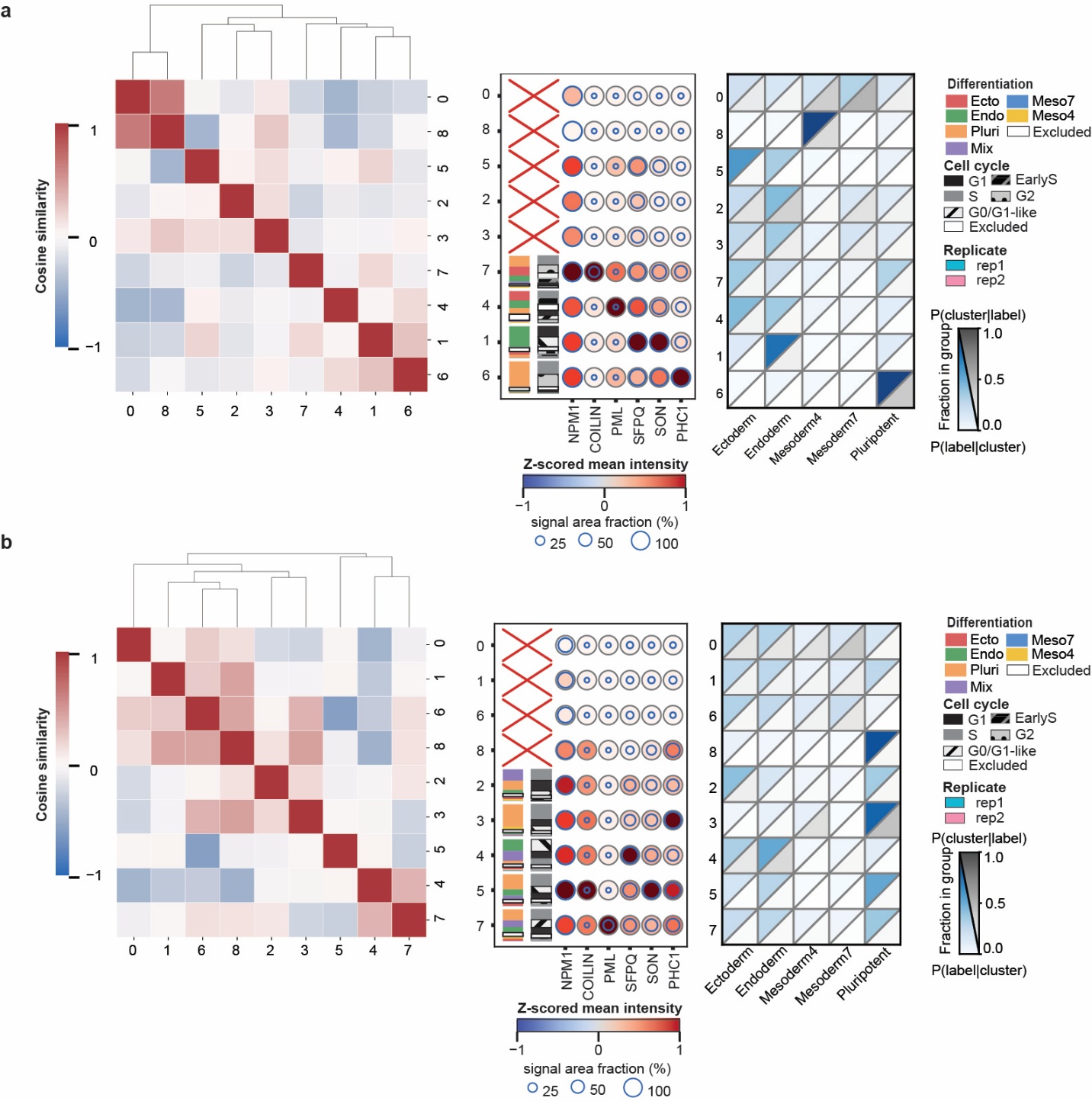

**Supplementary Fig. 11: Selection of organelle-scale clusters for finer subclustering.** **a,** Replicate 1. Left: cosine similarity between the latent-space profiles of organelle-scale Leiden clusters at resolution 0.8, ordered by hierarchical clustering. Middle: dot plot of MLO marker intensity per cluster, with side bars indicating germ layer/pluripotency group, cell-cycle state, and replicate. Clusters marked with a red cross were excluded from further sub-clustering, on the basis of high cosine similarity to another cluster combined with low overall MLO marker expression (see Methods). Right: bipartite correspondence between clusters and experimental differentiation groups (as in Supplementary Fig. 10a), used to assign germ layer/pluripotency identity to each cluster. **b,** As in a, for replicate 2.

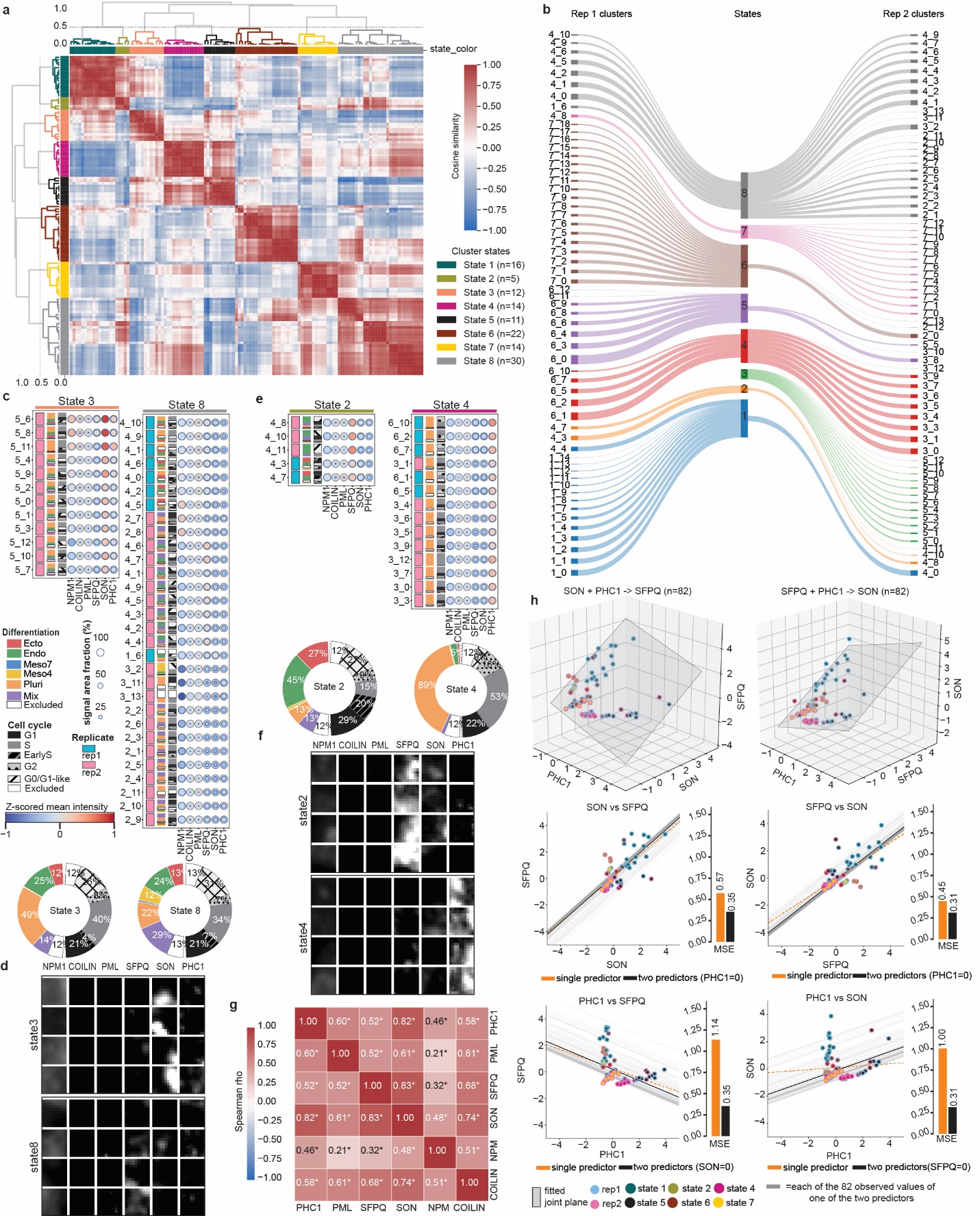

**Supplementary Fig. 12: Complete organelle-scale state clustering, marker profiles for every state, and extension of the joint linear model analysis to additional target markers.** **a,** Cosine similarity between all 124 organelle-scale subclusters, ordered by average-linkage hierarchical clustering. The colored bar indicates the 8 states obtained by cutting this dendrogram at a distance of 0.5 (state sizes as indicated). **b,** Sankey diagram showing the distribution of replicate 1 and replicate 2 patches across the 8 states; edge width is proportional to the percentage of each replicate's patches assigned to a given state. **c,** Dot plots and abundance-weighted composition donuts for states 3 and 8, the two states excluded from downstream cross-marker analyses. **d,** Representative organelle-scale MLO images for the excluded states shown in c **e,** Dot plots and abundance-weighted composition donuts for states 2 and 4, the two out of six states retained for downstream cross-marker analyses **f,** Representative organelle-scale MLO images for the retained states shown in e. **g,** Three-dimensional joint linear models predicting SFPQ and SON, extending the joint linear model partial-regression comparison in Fig. 4h to these two additional target markers (see Methods). Below, two-dimensional projections of each model compare the marginal regression line (“single predictors” to the joint model's partial-regression line (“two predictors.”, remaining marker fixed at 0), with mean squared error (MSE) of each model's predictions shown alongside. Points are colored by state and outlined by replicate. n = 82 subclusters.

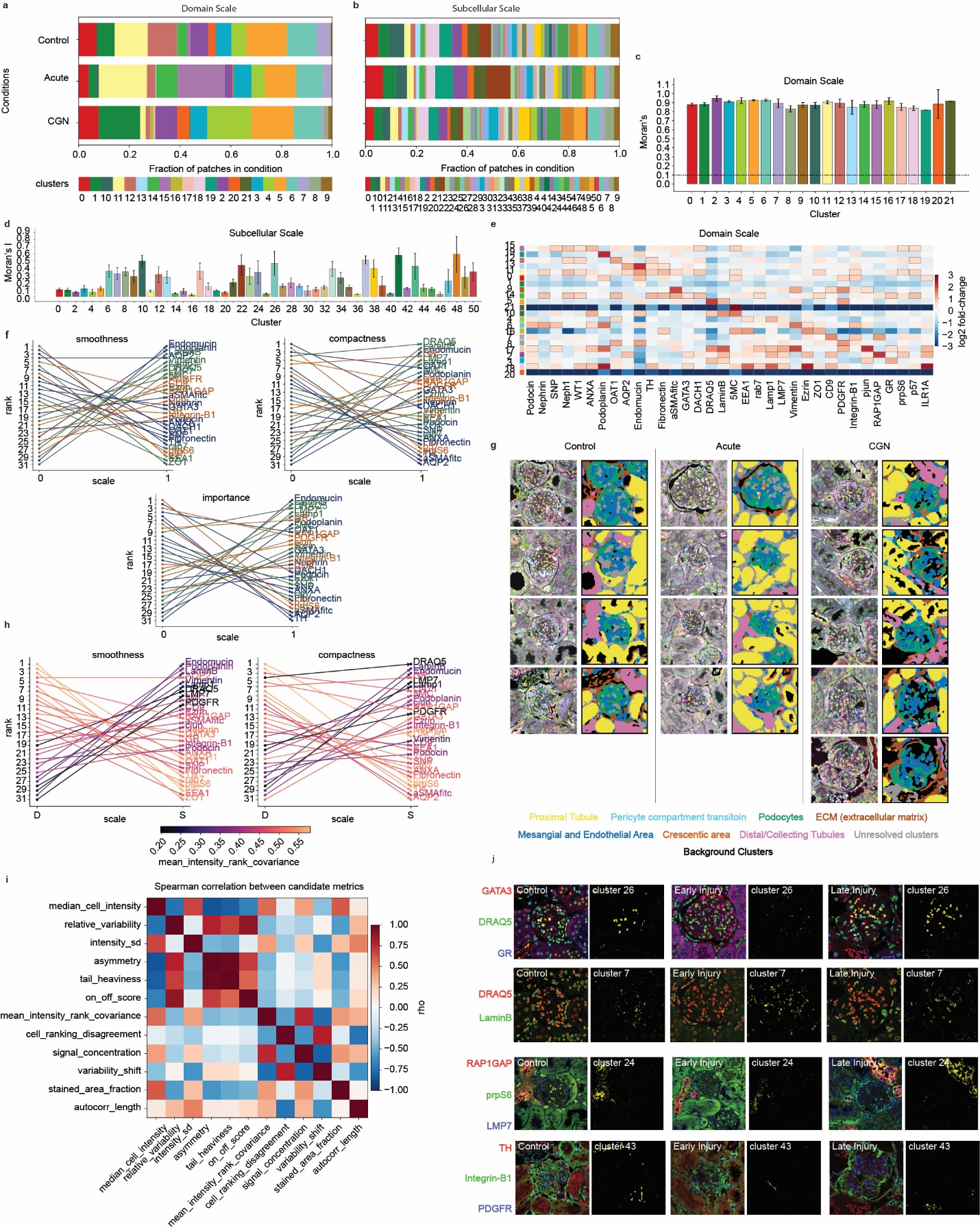

**Supplementary Fig. 13: CGN analysis of scaleVAE: a** cluster fraction distribution of the domain scale, **b** cluster fraction distribution of the subcellular scale, **c** the computed Moran’s I per cluster averaged over slides in the domain scale, **d** the computed Moran’s I per cluster averaged over the slides in the subcellular scale, **e** z-scored stain expression distribution over the clusters of the domain scale prior to domain mapping, **f**  the change in rank between scales colored by cellular, subcellular IDs and signaling pathways categories, **g** spatial projection of clusters of domain scale into individual domains across all validation set’s slides, **h** the change in rank between scales colored by the computed MIRC score for smoothness and compactness associated to the importance plot in Fig. 5h, **i** the spearman correlation of candidate metrics introduced in Fig. 5g, **j** example cluster projections of the subcellular scale with mesangial cells in cluster 26, pericyte-like cells in cluster 7, the mTOR signal localized in the distal/collecting tubules in cluster 24 and the sympathetic nerves in cluster 43.

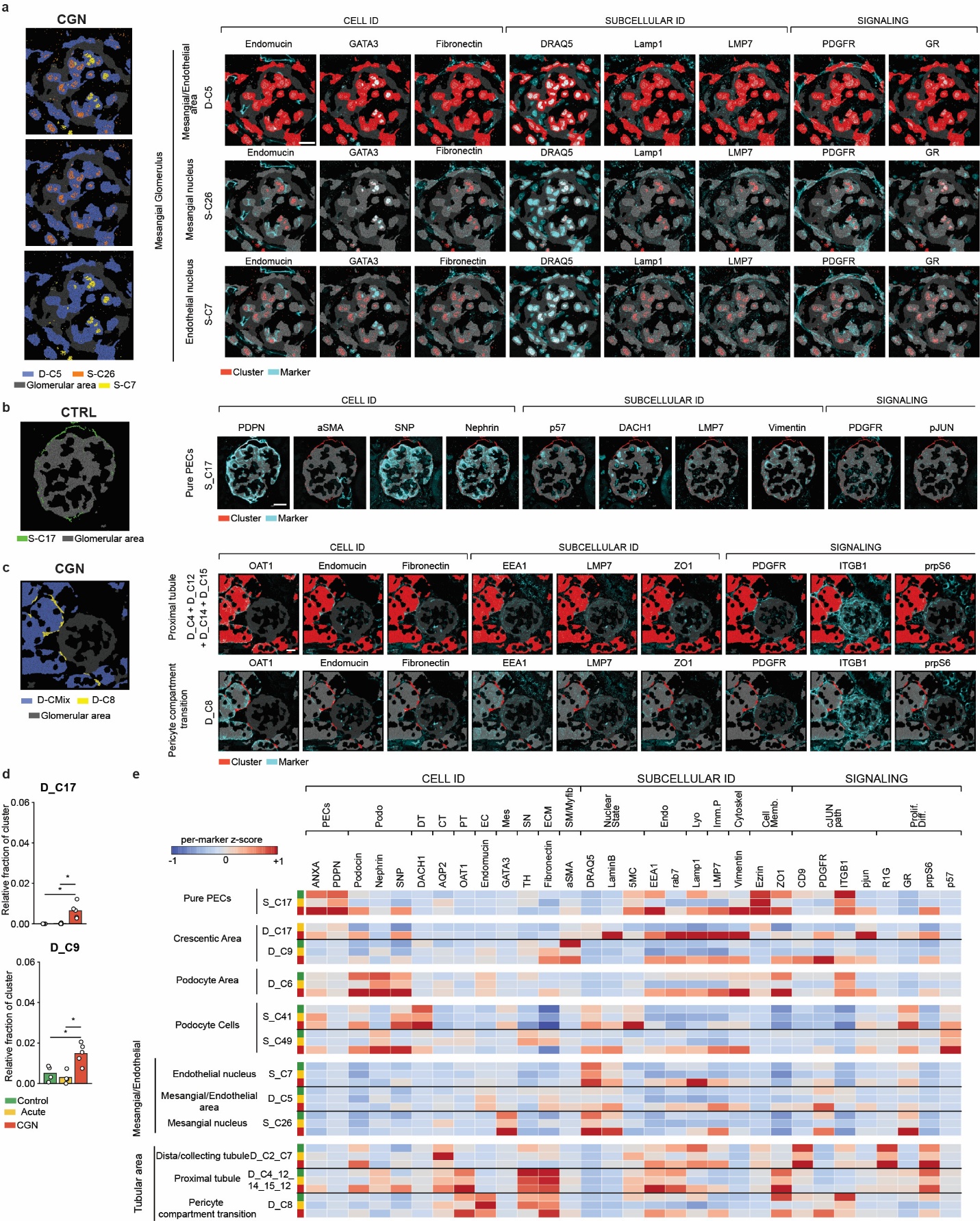

**Supplementary Fig. 14. Marker signatures of the mesangial, parietal and tubular compartments resolved by scaleVAE**. **a**, Left, cluster maps of the mesangial compartment on representative CGN glomeruli: D-C5 (blue), S-C26 (orange) and S-C7 (yellow) over the remaining glomerular area in grey. Right, each cluster mask (red) overlaid on one marker channel (cyan); white indicates pixels where the two coincide, grey the glomerular area behind. Rows, Mesangial/Endothelial area (D-C5), Mesangial nucleus (S-C26) and Endothelial nucleus (S-C7). Columns are grouped as CELL ID, SUBCELLULAR ID and SIGNALING. Scale bars, 200 px **b**, As in a, for S_C17 (Pure PECs, green) on a representative Control glomerulus, against ten markers. Scale bars, 200 px (16 µm) **c,** As in a, for the proximal tubule (D_C4 + D_C12 + D_C14 + D_C15, blue) and the pericyte compartment transition (D_C8, yellow) on a representative CGN section, against nine markers. Scale bars, 200 px **d**, Relative abundance of D_C17 and D_C9, given as the fraction of a sample's patches assigned to the cluster. Bars, group mean; dots, individual ROIs, one per kidney slide. Samples; n = 4 Control, n = 4 Acute and n = 5 CGN ROIs. Exact two-sided permutation test on the difference of means; * p < 0.05. **e**, Per-marker z-scored intensity for twelve cluster entries, grouped into Pure PECs, Crescentic Area, Podocyte Area, Podocyte Cells, Mesangial/Endothelial and Tubular area. Each entry carries one sub-row per condition, marked by the green (Control), yellow (Acute) and red (CGN) chip on the left; D_C17 has two, as it is absent from Control. Merged entries are averaged across their member clusters within each condition. Intensities were z-scored per marker across the 35 sub-rows shown, divided by the 99th percentile of |z| for that marker and clipped to ±1. **Abbreviations**: **Cell ID** — PECs, parietal epithelial cells (ANXA, Annexin A3; PDPN, podoplanin); Podo, podocytes (Podocin; NPHN, nephrin; SNP, synaptopodin); DT, distal tubule (DACH1); CT, collecting tubule/duct (AQP2, aquaporin-2); PT, proximal tubule (OAT1, organic anion transporter 1); EC, endothelial cells (EMCN, endomucin); Mes, mesangial cells (GATA3); SN, sympathetic nerve (TH, tyrosine hydroxylase); ECM, extracellular matrix (FN, fibronectin); SM/Myfib, smooth muscle/myofibroblast (aSMA, alpha-smooth muscle actin). **Subcellular ID** — Nuclear State (DRAQ5, DNA dye; LaminB, Lamin B1; 5MC, 5-methylcytosine); Endo, endosome (EEA1, early endosome antigen 1; rab7, Ras-associated binding protein 7); lyo, lysosome (Lamp1, lysosomal associated membrane protein 1); ImmP, immunoproteasome (LMP7); cytoskel, cytoskeleton (Vimentin); Cell Memb., cell membrane (Ezrin; ZO1, zonula occludens-1). **Signaling** — cJUN path (CD9; PDGFR, platelet-derived growth factor receptor beta; ITGB1, integrin beta 1; pJun, phospho-c-Jun); Prolif diff, proliferation/differentiation (R1G, RAP1GAP; GR, glucocorticoid receptor; prpS6, phospho-ribosomal protein S6; p57, CDKN1C).
